# Release of Notch activity coordinated by IL-1β signalling confers differentiation plasticity of airway progenitors via Fosl2 during alveolar regeneration

**DOI:** 10.1101/2021.06.19.449004

**Authors:** Jinwook Choi, Yu Jin Jang, Catherine Dabrowska, Elhadi Iich, Kelly V. Evans, Helen Hall, Sam M. Janes, Benjamin D. Simons, Bon-Kyoung Koo, Jonghwan Kim, Joo-Hyeon Lee

**Author notes:** First author (Jinwook Choi, Yu Jin Jang).

## Abstract

While the acquisition of cellular plasticity in adult stem cells is essential for rapid regeneration after tissue injury, little is known about the underlying molecular mechanisms governing this process. Our data reveal the coordination of airway progenitor differentiation plasticity by inflammatory signals during alveolar regeneration. Upon damage, IL-1β signalling-dependent modulation of Jag1/2 expression in ciliated cells results in the inhibition of Notch signalling in secretory cells, which drives reprogramming and acquisition of differentiation plasticity. We identify a core role for the transcription factor Fosl2/Fra2 in secretory cell fate conversion to alveolar type 2 (AT2) cells retaining the distinct genetic and epigenetic signatures of secretory lineages. We furthermore reveal that KDR/FLK-1^+^ human secretory cells display a conserved capacity to generate AT2 cells via Notch inhibition. Our results demonstrate the functional role of a IL-1β-Notch-Fosl2 axis for the fate decision of secretory cells during injury repair, proposing a new potential therapeutic target for human lung alveolar regeneration.

## Main

Tissue homeostasis is maintained by stem and progenitor cells capable of self-renewal and lineage-restricted differentiation, governed by specialised microenvironments. Both stem and progenitor cells acquire a remarkable potential for multi-lineage differentiation after tissue insults to allow for rapid regeneration of damaged tissue^1, 2^. These capacities depend on the nature and/or extent of injury, the repair ability of resident stem/progenitor cells, and their local microenvironments. Dysregulation of cellular plasticity, however, leads to lineage infidelity, and is implicated in various diseases^2^. Thus, to understand the process of tissue repair and regeneration in homeostasis and pathology, it is important to elucidate how stem and progenitor cells sense damage signals and control transient plasticity for fate conversion.

Lung tissue is maintained by several stem and progenitor cell types that reside in anatomically distinct regions alongside the pulmonary axis^3, 4^. In the distal lungs two main stem cell types, secretory cells and alveolar type 2 (AT2) cells, separately maintain the airway and alveolar epithelium compartments, respectively. In alveolar homeostasis and injury conditions, AT2 cells have been identified as the resident alveolar stem cells capable of self-renewal and differentiation into alveolar type 1 (AT1) cells^5–7^. However, when alveolar integrity is severely disrupted following lung injury, such as bleomycin or influenza infection, stem/progenitor cells localised in the distal airway (including secretory cells) can contribute to alveolar regeneration by differentiating into AT2 cells^5, 6, 8–12^. In particular, several lines of evidence reveal the capacity of lineage-traced *Scgb1a1^+^* airway secretory cells, which exhibit restricted capacity to generate airway cells in homeostasis, to produce new AT2 cells following severe lung injury in order to achieve effective repair of the alveolar epithelium^5, 6^. However, little is known about the cellular events and molecular cues which dictate how secretory cells lose their identity in response to alveolar damage and subsequently acquire AT2 cell fate. Significantly, the chronic loss of alveolar integrity is strongly associated with various human lung diseases. Thus, it is imperative to understand whether there is a functionally conserved secretory cell population that readily mediates human alveolar regeneration, and whether regulatory mechanisms are conserved between human and mouse lungs.

Here, we establish chemically defined feeder-free *in vitro* organoid cultures wherein secretory cells and AT2 cells are capable of expansion with a restricted differentiation capacity. This platform enables us to perform robust interrogation of the molecular and cellular behaviour of secretory and AT2 cells *in vitro*. By comparing the gene expression patterns in these organoids, we identify the Notch signalling pathway as a pivotal regulator for the differentiation plasticity of secretory cells into AT2 cells. We demonstrate the role of IL-1β-mediated inflammation in ciliated cells in regulating this differentiation plasticity *in vivo*. The AP-1 transcription factor Fosl2/Fra2 is defined as an essential transcription factor for AT2 cell conversion from secretory cells in response to Notch signalling. Further, the distinct genetic and epigenetic signatures of secretory-derived AT2 cells confer their functional difference compared to resident AT2 cells. Lastly, we demonstrate that the role of Notch signalling in the transition of secretory cells to AT2 cells is conserved in human airway organoids derived from KDR/FLK-1^+^ secretory cells. Our study illustrates the precise molecular and cellular coordination required for secretory cell-mediated alveolar regeneration, providing new insights into both the potential regenerative roles of these cells and regulatory networks in repairing alveolar destruction in lung disease.

## Results

### Establishment of *in vitro* feeder-free self-renewing culture conditions for distal stem cells

To better understand and more easily manipulate lung epithelial stem cells, we established a simple feeder-free organoid culture condition with defined factors which support the molecular and functional identity of stem cells over long-term culture. Airway secretory cells expressing *Scgb1a1* (*Secretoglobin 1a1*, also known as *CC10* or *CCSP*) were isolated from the distal lung tissues of secretory cell reporter mice (*Scgb1a1-CreER^TM/+^*;*R26R^tdTomato/+^*) after tamoxifen treatment^13^ (Fig. 1a). Lineage-labelled secretory cells were embedded in Matrigel supplemented with WNT3A, RSPO1 (R-spondin 1), EGF, FGF7, FGF10, and NOG (Noggin), factors that are known to support the growth of human embryonic lung tip cells and progenitors derived from foregut lineages^14–18^ (Fig. 1b). Under this condition, we were able to grow secretory cells that formed cyst-like organoids (Fig. 1b). The organoids were capable of long-term expansion (>2 years) by dissociation into single cells with stable forming efficiency (Fig. 1c,d). We also successfully generated organoids after limiting dilution with drops containing single cells (Fig. 1e), which serves as evidence of the clonogenicity and stem cell property of secretory cells maintained in this condition. FGF10 was dispensable for organoid growth with no significant difference in the efficiency of organoid formation and maintenance over serial passages (Fig. 1b-d). Without FGF7, however, secretory cells failed to form organoids (Fig. 1b,d). Furthermore, the absence of factors supporting WNT activity resulted in deterioration of organoid expansion over passages (Fig. 1b,d), indicating that both FGF7 and WNT signalling are essential for the self-renewal of distal secretory cells. The initial organoid cultures retained heterogeneous phenotypes of organoids, including airway retaining CC10^+^ secretory and Acetylated-Tubulin (Act-Tub)^+^ ciliated cells, alveolar retaining SPC^+^ AT2 and HOPX^+^ AT1 cells, and mixed organoids retaining both airway and alveolar lineage cells, similar to previous findings from the co-culture system^19^ (Fig. 1f,i). After five passages, organoids expanded to form mainly mixed organoids with budding-like morphology over multiple passages (Fig. 1g- i).

**Figure 1.**
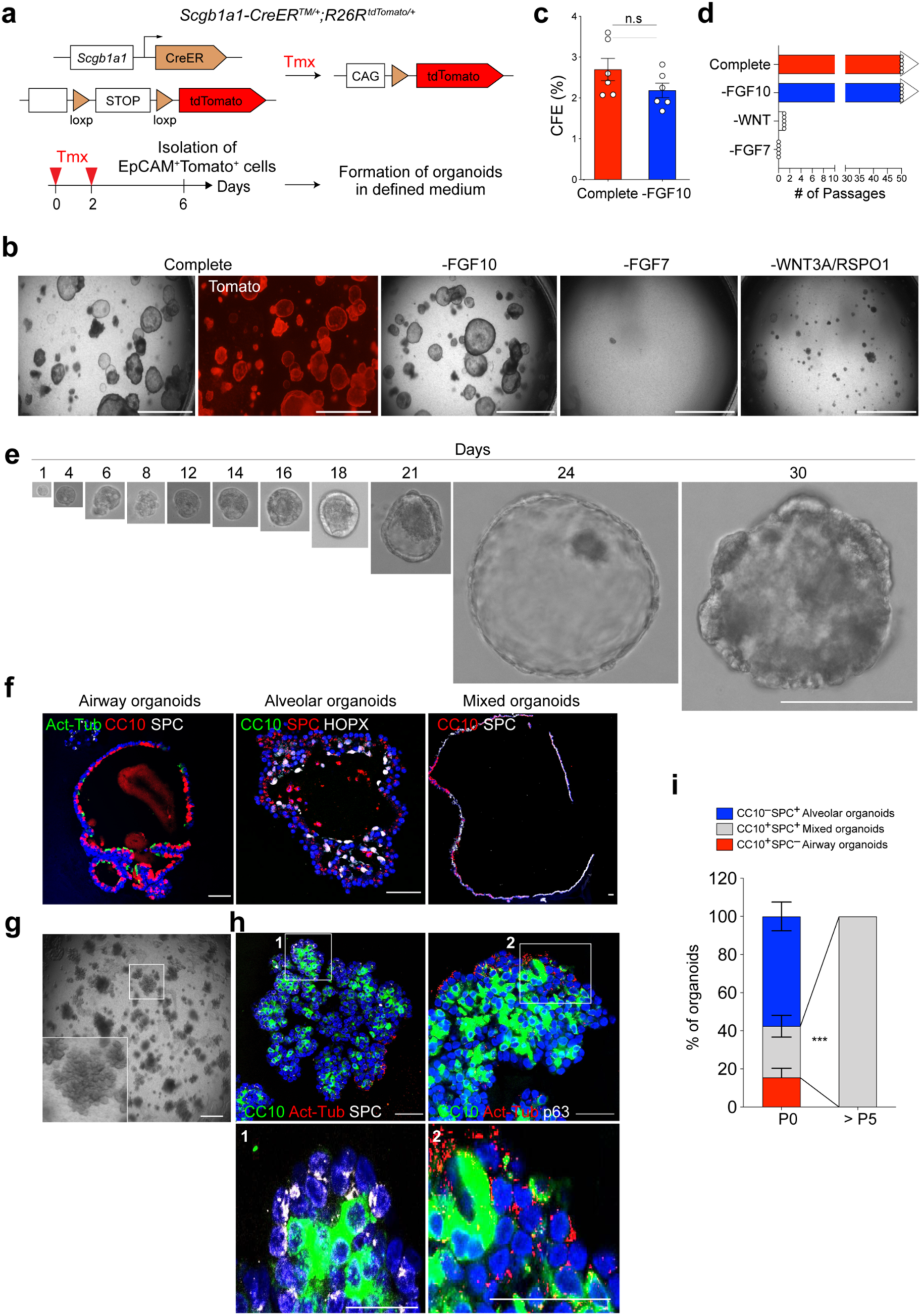
Establishment of feeder-free organoids derived from distal secretory cells. **a,** Experimental design for isolation of *Scgb1a1* lineage-labelled cells. *Scgb1a1* lineage- labelled cells were isolated at day 4 after final tamoxifen treatment. **b,** Representative bright- field or fluorescent images of organoids derived from lineage-labelled *Tomato^+^Scgb1a1^+^* cells in indicated conditions; complete medium (See **Methods**) with WNT3A, RSPO1 (R-spondin 1), EGF, FGF7, FGF10, and NOG (Noggin), withdrawal of indicated factors (−FGF10, −FGF7, or −WNT3A/RSPO1). Scale bar, 2,000µm. **c, d,** Statistical quantification of colony forming efficiency (**c**) and passaging numbers (**d**) of organoids. Each individual dot represents one biological replicate and data are presented as mean and s.e.m. n.s; not significant. **e,** Representative serial bright-field images of a lung organoid growing originated from single *Tomato^+^Scgb1a1^+^* cells at the indicated time points. Magnifications: X20 (day 0 and 4), X10 (day4-21), and X4 (day 24 and 30). Scale bars, 200µm. **f,** Representative immunofluorescence (IF) images of three distinctive types of organoids derived from *Tomato^+^Scgb1a1^+^* cells at the establishment under feeder-free condition with complete culture medium; Airway organoids retaining CC10^+^ secretory and Act-Tub^+^ ciliated cells (left), Alveolar organoids retaining SPC^+^ AT2 and HOPX^+^ AT1 cells (middle), and Mixed organoids retaining both CC10^+^ secretory and SPC^+^ AT2 cells (right). CC10 (for secretory cells, red or green), Acetylated-Tubulin (Act-Tub, for ciliated cells, green), SPC (for AT2 cells, white), HOPX (for AT1 cells, white), and DAPI (blue). Scale bars, 50µm. **g,** Representative bright-field images of organoids under feeder-free condition with complete culture medium at passage 5. Insets show high-power view. Scale bars, 2,000µm. **h,** Representative IF images of mixed organoids cultured in complete medium at passage 5. CC10 (green), Act-Tub (red), SPC (white, left), p63 (for basal cells, white, right), and DAPI (blue). Insets (1, 2) show high-power view. Scale bars, 50µm. **i,** Quantification of each organoid types in complete medium at passage 0 and over passage 5; Airway organoids (CC10^+^SPC^−^; red), Alveolar organoids (CC10^−^SPC^+^; blue), and Mixed organoids (CC10^+^SPC^+^; grey). Data are presented as mean and s.e.m. (n=3 independent experiment). ***p<0.001.

We next asked whether our culture condition could allow AT2 cells to form organoids. Lineage-labelled AT2 cells were isolated from *Sftpc-CreER^T2/+^*;*R26R^tdTomato/+^* reporter mice after tamoxifen treatment^5^ (Extended Data Fig. 1a). Like secretory cell-derived organoids (referred to here as SCOs), they were capable of forming cyst-like organoids (Extended Data Fig. 1b). Consistent with previous studies^20–23^, FGF7 and WNT activity were essential for forming and maintaining AT2 cell-derived organoids (referred to hereafter as ACOs), which contained both SPC^+^ AT2 and Pdpn^+^ AT1 cells (Extended Data Fig. 1b-f). We also confirmed the clonogenic ability of AT2 cells by organoid culture with single cells (Extended Data Fig. 1e). However, unlike SCOs, which tolerate the presence of FGF10 in long-term culture, ACOs were better maintained in the absence of FGF10 (Extended Data Fig. 1d). After several passages, these organoids expanded to form alveolar-like structures that were mainly composed of AT2 cells with little AT1 cell differentiation (Extended Data Fig. 1g,h). Taken together, we developed a feeder-free organoid culture system with chemically defined medium, wherein distal airway secretory cells and AT2 cells were maintained with stem cell activity in long-term cultures. Activation of both FGF7 and WNT signalling pathways was sufficient to support them.

### Inhibition of Notch signalling confers plasticity of secretory cells to differentiate into AT2 cells upon injury

To investigate the characteristics of SCOs and ACOs, we first performed bulk RNA-seq analysis. Consistent with immunofluorescent (IF) staining data, SCOs exhibited a distinct gene expression pattern enriched in secretory cell markers such as *Scgb1a1*, *Cyp2f2*, and *Gabrp*, while AT2 cell markers including *Sftpc* and *Etv5* were highly expressed in ACOs (Fig. 2a). In addition, gene set enrichment analysis (GSEA) revealed that secretory cell marker genes were enriched in the genes highly active in SCOs, whereas AT2 cell markers were enriched in ACOs (Fig. 2b). Notably, GSEA indicated that Notch signalling was distinctly enriched in SCOs (Fig. 2b). We also confirmed that the expression of genes regulated by Notch signalling, such as *Hes1*, *Hes6* and *Nrarp*, was upregulated in SCOs compared to ACOs (Fig. 2a). Additional IF staining also confirmed the expression of intracellular domain of Notch1 (N1ICD) in SCOs (Extended Data Fig. 2a). This observation prompted us to examine whether Notch signalling is involved in the identity and fate behaviour of secretory cells. We dissociated established SCOs into single cells and replated them to form organoids in the presence of DMSO (control) or DAPT, a γ-secretase inhibitor, to inhibit Notch signalling. Remarkably, SPC^+^ AT2 cells were dramatically increased at the expense of CC10^+^ secretory cells in DAPT-treated organoids compared to control, whereas other lineages were barely affected (Fig. 2c). To further characterise these organoids, we performed RNA-seq analysis of SCOs with and without DAPT treatment. Unlike control organoids, DAPT-treated organoids displayed strongly upregulated AT2 cell marker gene expression and a substantial reduction of secretory cell markers (Fig. 2d). GSEA also showed similar expression patterns (Fig. 2e). To directly confirm the functional effect of Notch activity in secretory cells, we knocked-down *Rbpj*, the major transcriptional effector of Notch signalling, in SCOs. Knockdown (KD) of *Rbpj* promoted the differentiation of secretory cells into AT2 cells without DAPT treatment (Extended Data Fig. 2b,c), suggesting that inhibition of Notch signalling triggers the conversion of secretory cells into AT2 lineage.

**Figure 2.**
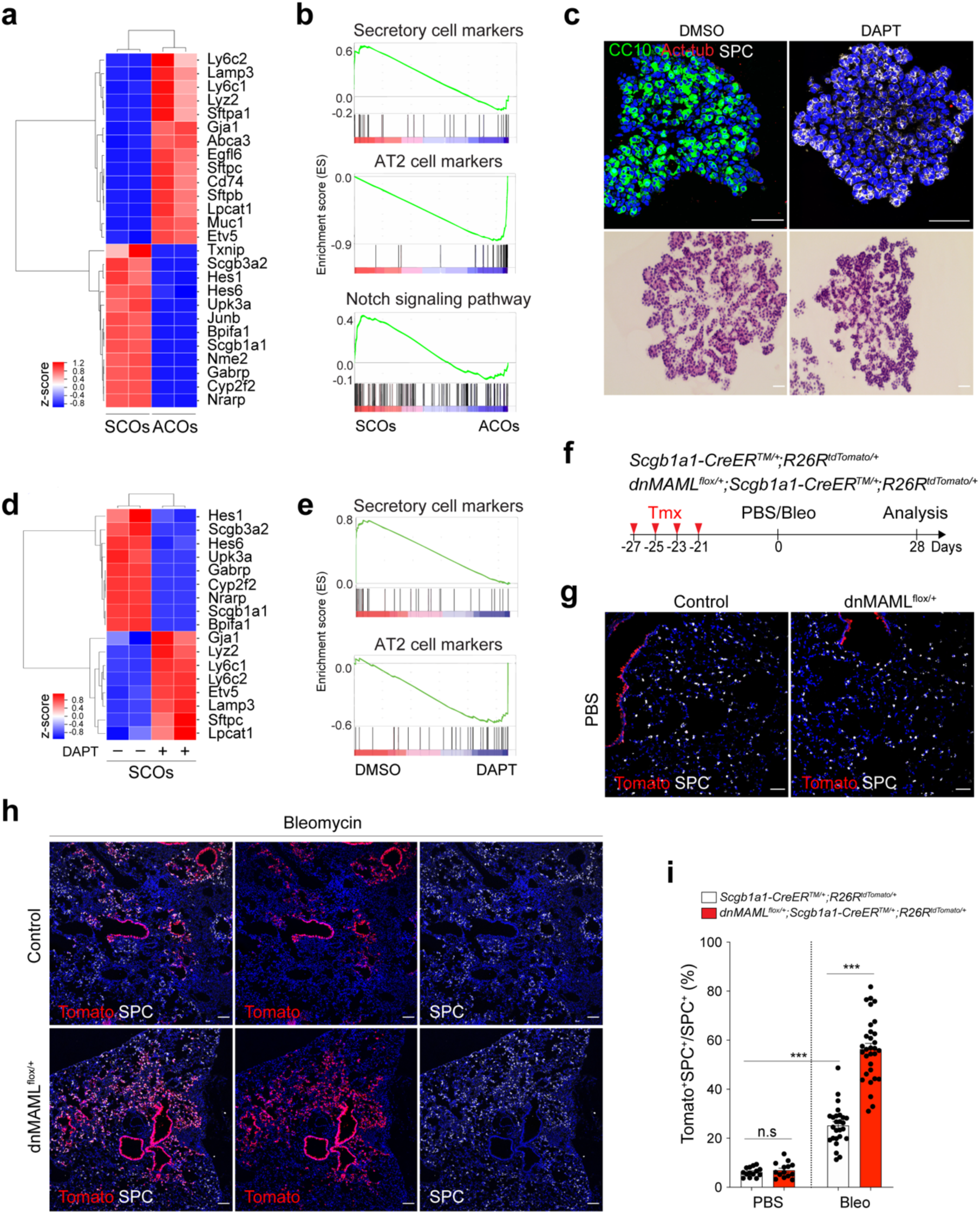
Enhanced differentiation of secretory cells into AT2 cells by inhibition of Notch activity upon lung injury. **a,** A heatmap showing normalised expression data for secretory and AT2 cell markers that were differentially expressed in secretory cell-derived organoids (SCOs) or AT2 cell-derived organoids (ACOs) cultured with defined media at passage 10. Values are z-scores. **b,** Gene set enrichment analysis (GSEA) showing the gene activity of the three gene sets for secretory cells (top), AT2 cells (middle), and Notch signalling pathway-related in SCOs or ACOs **c,** Representative IF (top) and H&E (bottom) images of organoids in treatment with DMSO (control) and DAPT at day 14. DAPT (20µM) were treated every other day during organoid culture. CC10 (green), Acetylated-Tubulin (Act-Tub, red), and SPC (white), and DAPI (blue). Scale bar, 50µm. **d,** A heatmap showing normalised expression data for Notch signalling related, secretory and AT2 cell marker genes that were differentially expressed in SCOs with or without DAPT treatment for 14 days. Values are z-scores. **e,** GSEA with two gene sets representing markers for secretory (left) and AT2 cells (right) in SCOs with or without DAPT treatment. **f,** Experimental design of lineage-tracing analysis for contribution of secretory cells into alveolar lineages by inhibition of Notch signalling using *Scgb1a1- CreER^TM/+^*;*R26R^tdTomato/+^* and *dnMAML^flox/+^*;*Scgb1a1-CreER^TM/+^*;*R26R^tdTomato/+^* mice after bleomycin (Bleo) injury. Specific time points for tamoxifen injection and tissue analysis are indicated. **g, h,** Representative IF images showing the derivation of *Scgb1a1* lineage-labelled AT2 cells in PBS-treated control (**g**) or bleomycin-treated (**h**) mice at day 28. Tomato (for *Scgb1a1* lineage, red), SPC (white), and DAPI (blue). Scale bar, 100µm. **i,** Statistical quantification of *Scgb1a1^+^* lineage-labelled AT2 cells at day 28 post PBS or bleomycin treatment. Each individual dot represents one section and data are presented as mean ± s.e.m. (n=2 biological replicates for PBS control, n=4 for bleomycin). ***p<0.001, n.s; not significant.

To address this finding *in vivo*, we used mice carrying a conditional dominant-negative mutant of mastermind-like 1 (dnMAML^flox/flox^)^24^, which inhibits NICD-induced Notch signalling, crossed with *Scgb1a1-CreER^TM/+^*;*R26R^tdTomato/+^*. Control (*Scgb1a1-CreER^TM/+^*;*R26R^tdTomato/+^*) or dnMAML^flox/+^ (*dnMAML^flox/+^*;*Scgb1a1-CreER^TM/+^*;*R26R^tdTomato/+^*) mice were exposed to PBS or bleomycin after tamoxifen treatment to induce alveolar injury (Fig. 2f). Inhibition of Notch activity by *dnMAML* expression was confirmed by assessing the expression of Hes1, a potent target gene of Notch signalling (Extended Data Fig. 2d,e). Consistent with previous studies^25–27^, the proportion of lineage-labelled ciliated cells was increased by Notch inhibition in dnMAML^flox/+^ compared to control mouse lungs (Extended Data Fig. 2f,g). As reported^13^, 6.23±1.09% of lineage-labelled AT2 cells were observed in the alveolar region of uninjured control lung (Fig. 2g,i). The percentage of Tomato^+^AT2 cells (6.97±13.12%) in uninjured lung of dnMAML^flox/+^ mice was similar to that seen in control mice (Fig. 2g,i). However, bleomycin injury significantly increased the population of lineage-labelled AT2 cells to 25.24±8.08% in control mice (Fig. 2h,i). Importantly, we observed a substantial increase in lineage-labelled AT2 cells (56.39±12.91%) post injury in dnMAML^flox/+^ mice (Fig. 2h,i), suggesting that persistent inhibition of Notch signalling in secretory cells enhances their differentiation towards AT2 cell fate during injury repair. Further analysis of *in vitro* organoids also demonstrated that *dnMAML*-expressing secretory cells enhanced the frequency of formation of alveolar organoids retaining AT2 cells and reduced occurrence of airway organoids retaining secretory cells (Extended Data Fig. 3). Moreover, pharmacological inhibition of Notch activity by DAPT treatment *in vivo* also showed a significant increase of AT2 cells derived from secretory cells (Extended Data Fig. 4).

Given that *Scgb1a1^+^* lineage-labelled cells contained heterogeneous populations, including CC10^+^SPC^+^ dual-positive cells, we further evaluated the differentiation capacity of lineage-labelled secretory cells into AT2 lineages via Notch inhibition after excluding lineage-labelled SPC^+^ AT2 cells. We isolated lineage-labelled secretory cells (EpCAM^+^Tomato^+^MHCII^−^) from control and dnMAML^flox/+^ mouse lungs, followed by transplantation into the lungs after bleomycin injury (Extended Data Fig. 5a,b). Secretory cells isolated from dnMAML^flox/+^ lungs showed enhanced ability to generate AT2 cells compared to control secretory cells (Extended Data Fig. 5c-h). We also confirmed that *dnMAML*-expressing secretory cells revealed the greater frequency of alveolar organoid formation compared to control secretory cells (Extended Data Fig. 5i). Taken together, our data strongly suggest that persistent inhibition of Notch signalling enhances the differentiation of secretory cells into AT2 cells post injury.

### Sustained activation of Notch signalling impairs the differentiation of secretory cells into AT2 cell fate during alveolar injury repair

We then asked whether the constitutive activation of Notch signalling affects the cell fate of secretory cells upon injury. To this end, we used *Red2-Notch^N1ICD^* mice, where constitutive *N1ICD* is specifically co-expressed in *tdimer2 red fluorescent protein* (RFP)^+^ cells in the original confetti mouse line (Yum *et al.*, in press) (Fig. 3a). The other three lineage outcomes, namely YFP, GFP and CFP, all represent wild-type events. In *in vitro* organoid cultures of YFP^+^ (wild-type, control) and RFP^+^ (constitutive *N1ICD*) cells, RFP^+^ cells showed a significant increase in organoid formation efficiency compared to that of YFP^+^ cells (Fig. 3b,c). Consistent with previous results (Extended Data Fig. 3), YFP^+^ control cells gave rise to three distinct types of organoids in co-cultures (Fig. 3d,e). However, only airway organoids arose from RFP^+^ cells (Fig. 3d,e). To further investigate the cellular behaviour of secretory cells with constitutive Notch activity *in vivo*, *Scgb1a1-CreER^TM/+^*;*Red2-Notch^N1ICD/+^* mice were administered PBS or bleomycin injury (Fig. 3a). In uninjured lungs, YFP^+^ and RFP^+^ cells were localised in airways with the expression of the secretory cell marker CC10, and no lineage-labelled cells were observed in the alveolar region (Fig. 3f,h and Extended Data Fig. 5j). After bleomycin injury, we observed an increased frequency of YFP^+^ lineage-labelled AT2 cells expressing SPC in the alveolar regions (Fig. 3g-i). However, an increase of RFP^+^ lineage- labelled AT2 cells was significantly compromised after injury (Fig. 3g-i). Although RFP^+^ cells also expanded post injury, they still maintained identity of secretory cells retaining CC10 expression (Fig. 3g-i).

**Figure 3.**
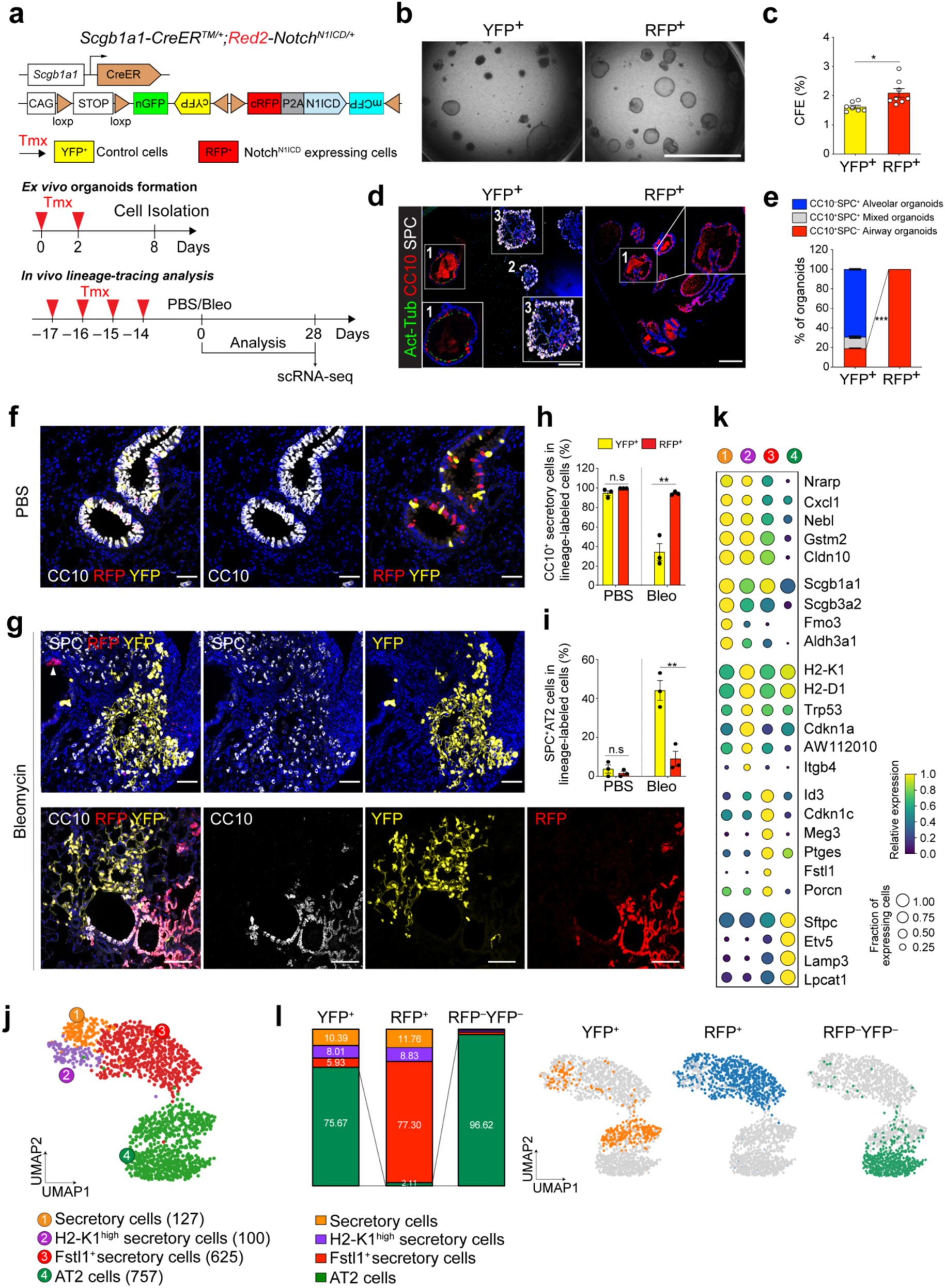
Impaired contribution of secretory cells into AT2 cell regeneration by sustained Notch activity. **a,** Experimental design of *ex vivo* organoid co-culture with stromal cells, lineage-tracing analysis, and single-cell RNA-sequencing (scRNA-seq) analysis using *Scgb1a1-CreER^TM/+^; Red2-Notch^N1ICD/+^* mice after bleomycin injury. Specific time points for tamoxifen injection and analysis for tissue and scRNA-seq are indicated. **b,** Representative bright-field images of organoids derived from control YFP^+^ (left) and *N1ICD*-expresing RFP^+^ (right) cells. For organoids, 5,000 cells of YFP^+^ or RFP^+^ cells were co-cultured with stromal cells with 1:5 ratio for 14 days. Scale bar, 2,000µm. **c,** Statistical quantification of colony forming efficiency (CFE) of organoids in (**b**). Each individual dot represents one individual biological replicate and data are presented as mean and s.e.m. *p<0.05. **d,** Representative IF images of three distinctive types of organoids in (**b**); Airway organoids (denoted as 1), Alveolar organoids (denoted as 2), and Mixed organoids (denoted as 3). CC10 (for secretory cells, red), Act-Tub (for ciliated cells, green), SPC (for AT2 cells, white), and DAPI (blue). Insets (1 and 3) show high-power view. Scale bars, 50µm. **e,** Quantification of each organoid types in (**d**); Airway organoids (CC10^+^SPC^−^; red), Alveolar organoids (CC10^−^SPC^+^; blue), and Mixed organoids (CC10^+^SPC^+^; grey). Data are presented as mean and s.e.m (n=3 biological replicates). ***p<0.001. **f,** Representative IF images showing the derivation of YFP^+^ or RFP^+^ cells in control mice at day 28 post PBS treatment. YFP (yellow), RFP (red), CC10 (white), and DAPI (blue). Scale bar, 50µm. **g,** Representative IF images showing the derivation of YFP^+^ or RFP^+^ cells at day 28 post bleomycin treatment. YFP (yellow), RFP (red), SPC (top, white), CC10 (bottom, white), and DAPI (blue). Arrowhead points to RFP^+^ cells. Scale bar, 50µm. **h, i,** Statistical quantification of CC10^+^ secretory (**h**) and SPC^+^ AT2 (**i**) cells derived from YFP^+^ or RFP^+^ cells at day 28 post PBS or bleomycin treatment. **p<0.01. **j,** Secretory and AT2 cell population were further analysed from scRNA-seq results (Extended Data Fig. 7b). Clusters of YFP^+^ and RFP^+^ lineage-labelled or YFP^−^RFP^−^ non-lineage labelled epithelial cells (1,609) from 10x Genomics 3’ single-cell RNA sequencing (scRNA-seq) analysis visualised by UMAP, assigned by specific colours. Number of cells in the individual cluster is depicted in the figure. **k,** Gene expression of key markers in each distinctive cluster. The size of the circle represents the fraction of cells expressing the gene and the colour represents the relative expression of each gene. **l,** Quantification (left) and UMAP (right) of distribution of each cluster across indicated lineage-labelled cells after injury.

We next sought to define the molecular identity and characteristics of YFP^+^ or RFP^+^ cells during alveolar regeneration. To this end, we carried out single-cell RNA-sequencing (scRNA-seq) analysis by isolating lineage-labelled (YFP^+^ or RFP^+^) and non-lineage-labelled (YFP^−^ RFP^−^) populations at day 28 post bleomycin injury (Extended Data Fig. 6a). Based on the expression of canonical markers for specific cell types, we revealed 14 distinct populations (Extended Data Fig. 6b,c). To gain insight into the impact of Notch activity on lineage differentiation of secretory cells into AT2 cells, we further analysed the secretory and AT2 cell populations (Fig. 3j). This analysis uncovered four distinctive clusters, including three different types of secretory cells and an AT2 cell cluster (Fig. 3j,k). In addition to the known secretory cell populations, including *Scgb3a2^high^* canonical secretory cells (cluster 1) and *H2-K1^high^* cells (cluster 2)^8^ sharing common secretory markers (e.g. *Cldn10*), our analysis identified an uncharacterized population which we termed *Fstl1^+^* secretory cells (cluster 3; Fig. 3j,k). While cells in this cluster still retained the identity of secretory cells based on *Scgb1a1* expression, they showed lower expression levels of some secretory cell markers such as *Scgb3a2* and *Cldn10* (Fig. 3k). Moreover, this cluster was marked by specific expression of discrete markers such as *Id3* and *Porcn* (Fig. 3k). Interestingly, this population had elevated expression of *Cdkn1c*/p57, involved in cell cycle inhibition, which could indicate the quiescent characteristics of this cluster *in vivo* (Fig. 3k and Extended Data Fig. 6d,e). Distribution of each cluster across collected samples allowed us to assess how control (YFP^+^) or *N1ICD*-expressing (RFP^+^) cells respond to injury. Consistent with lineage-tracing analysis, control YFP^+^ cells contributed to alveolar regeneration by giving rise to AT2 cells post injury (Fig. 3l). However, RFP^+^ cells were arrested at the stage of the *Fstl1*^+^ population, causing impaired differentiation into AT2 cells (Fig. 3l). Pseudotime analysis revealed that the transition from secretory cells to AT2 cells is mediated via this cluster 3, suggesting that this population may function as a transitional intermediate in the lineage transition between secretory cells and AT2 cells, and inactivation of Notch activity is required for the transition of this intermediate state into AT2 cell fate (Extended Data Fig. 6f). Overall, our results indicate that sustained activation of Notch signalling causes defects in alveolar regeneration by blocking the fate conversion from secretory cells to AT2 cells.

### IL-1β signalling in ciliated cells is required for Notch-dependent fate conversion of secretory cells to AT2 cells after injury

We next asked how Notch activity in secretory cells is regulated during regeneration. Given that Notch signalling involves short-range cellular communication through direct contact by receptor-ligand interaction, we examined which cells highly express the Notch ligands Jagged 1 (Jag1) and Jagged 2 (Jag2). Our scRNA-seq analysis from *Scgb1a1-CreER^TM/+^*;*Red2- Notch^N1ICD/+^* mice revealed that *Jag1* and *Jag2* are highly expressed in ciliated cells, which reside adjacent to secretory cells in the airway, consistent with previous studies^27–29^ (Extended Data Fig. 7a,b). Immunohistochemistry (IHC) staining also confirmed the high expression of Jag1 in ciliated cells (Extended Data Fig. 7c). Notably, we found that Jag1 expression was downregulated after bleomycin treatment, suggesting dynamic modulation of Notch signalling by ciliated cells during injury repair (Extended Data Fig. 7c). Previously we showed dynamic expression of IL-1β during regeneration after bleomycin injury^30^. Notably, *Il1r1*, a functional receptor for IL-1β, is specifically expressed in ciliated cells, in addition to subsets of AT2 cells (Extended Data Fig. 7d,e)^30^. Thus, we asked whether IL-1β signalling modulates the expression of Notch ligands in ciliated cells. The expression of Notch ligands, *Jag1* and *Jag2*, was significantly reduced in *Il1r1^+^* ciliated cells (EpCAM^+^*Il1r1*^+^CD24^high^) after bleomycin injury compared to PBS control (Fig. 4a,b). Furthermore, we treated PBS or IL-1β in isolated *Il1r1^+^* ciliated cells for 24 hours. The expression of *Jag1* and *Jag2* was significantly downregulated in IL-1β-treated ciliated cells compared to control cells, suggesting that IL-1β signalling directly modulates the expression of Notch ligands in ciliated cells after injury (Fig. 4c).

**Figure 4.**
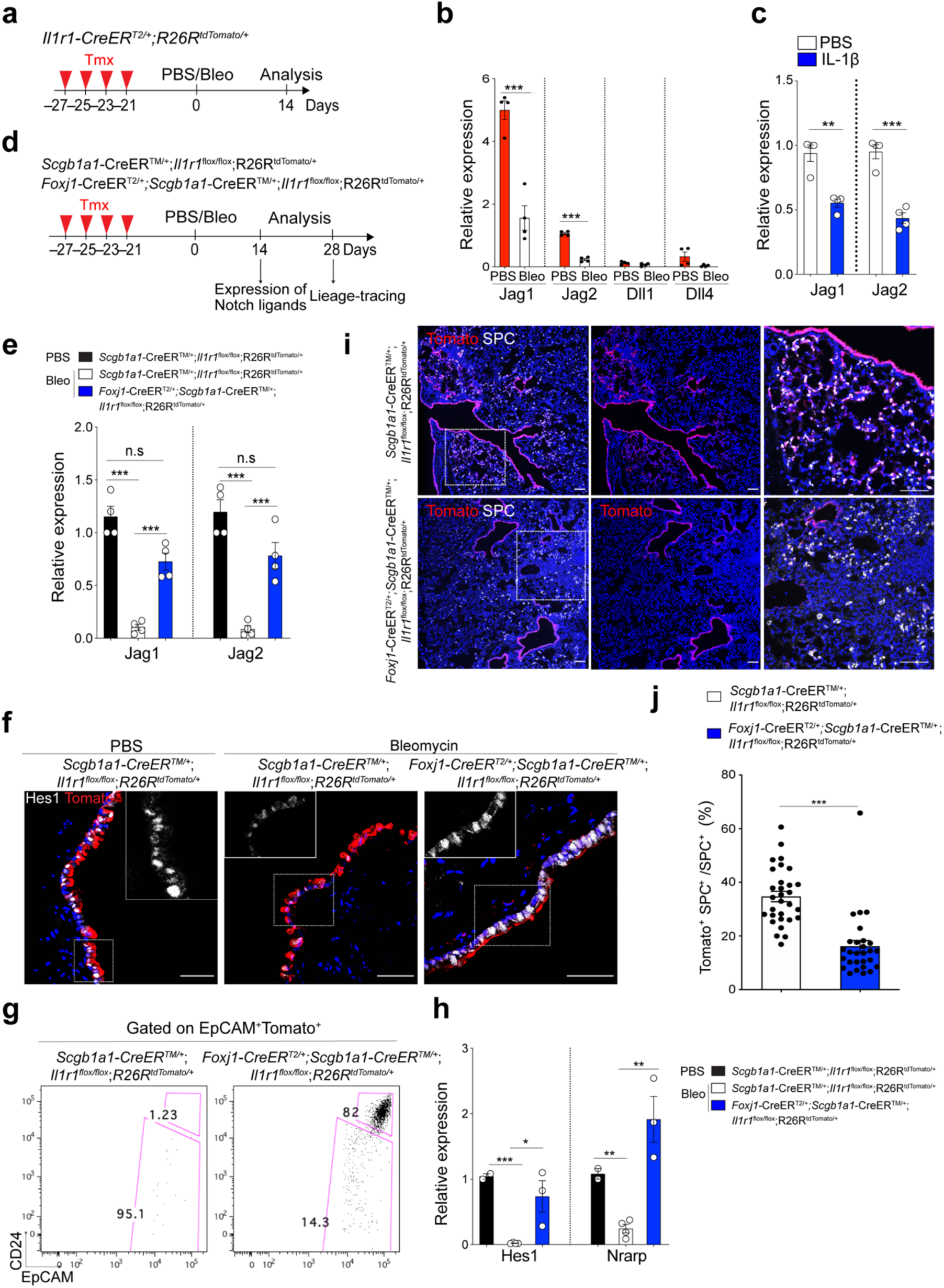
Regulation of Notch ligand expression in ciliated cells by IL-1β signalling. **a,** Experimental design for isolation of *Il1r1*-expressing ciliated cells after PBS or bleomycin treatment using *Il1r1-CreER^T2/+^*;*R26R^tdTomato/+^* mice. Specific time points for tamoxifen injection are indicated and tissue was analysed at day 14 post PBS or bleomycin treatment. **b,** Quantitative PCR (qPCR) analysis showing the expression of Notch ligands in isolated lineage- labelled ciliated cells (EpCAM^+^*Il1r1^+^*CD24^high^). ***p<0.001. **c,** qPCR analysis showing the expression of *Jag1* and *Jag2* in cultured ciliated cells treated with PBS or IL-1β. Isolated ciliated cells (EpCAM^+^*Il1r1^+^*CD24^high^) from *Il1r1-CreER^T2/+^*;*R26R^tdTomato/+^* mouse lungs after two doses of tamoxifen were cultured in Matrigel for 24 hrs in the presence of PBS or IL-1β (10ng/ml) *in vitro*. **p<0.01, ***p<0.001. **d,** Experimental design for lineage-tracing analysis of secretory cells after PBS or bleomycin treatment using *Scgb1a1- CreER^TM/+^*;*Il1r1^flox/flox^*;*R26R^tdTomato/+^* or *Foxj1-CreER^T2^*;*Scgb1a1-CreER^TM/+^*;*Il1r1^flox/flox^*; *R26R^tdTomato/+^* mice. Specific time points for tamoxifen injection and tissue analysis are indicated. **e,** qPCR analysis showing the expression of *Jag1* and *Jag2* in isolated ciliated cells (Tom^+^EpCAM*^+^*CD24^high^) after treatment of PBS or bleomycin. ***p<0.001, n.s; not significant. **f,** Representative IF images showing Hes1 expression in lineage-labelled secretory cells after bleomycin at day 14 post injury in the indicated genotype: Tomato (for *Scgb1a1* lineage, red), Hes1 (white), and DAPI (blue). Insets show high-power view of Hes1 expression. Scale bars, 50µm. **g,** Flow cytometry sorting strategy for lineage-labelled secretory cells after PBS or bleomycin injury. EpCAM^+^CD24^−^ cells gated from EpCAM^+^Tomato^+^ were used for isolation of lineage-labelled secretory cells and qPCR analysis. **h,** qPCR analysis of downstream target genes of Notch signalling including *Hes1* and *Nrarp* from isolated lineage- labelled secretory cells in (**g**). *p<0.05, **p<0.01, **p<0.001. **i,** Representative IF images showing the derivation of *Scgb1a1^+^* lineage-labelled AT2 cells in PBS-treated control or bleomycin-treated mice at day 28. Tomato (for *Scgb1a1* lineage, red), SPC (white), and DAPI (blue). White boxed insets show high-power view (right panel). Scale bar, 100µm. **j,** Statistical quantification of Tomato^+^SPC^+^ AT2 cells at day 28 post bleomycin treatment in (**f**). Each individual dot represents one section from four (*Scgb1a1-CreER^TM/+^*;*Il1r1^flox/flox^*;*R26R^tdTomato/+^*) or three (*Foxj1-CreER^T2^*;*Scgb1a1-CreER^TM/+^*;*Il1r1^flox/flox^*;*R26R^tdTomato/+^*) biological replicates. Data are presented as mean ± s.e.m. ***p<0.001.

We next asked whether depletion of IL-1β signalling on ciliated cells impacts the fate behaviour of secretory cells after injury *in vivo*. To delete *Il1r1* in ciliated cells and lineage-trace secretory cells simultaneously, we established *Il1r1^flox/flox^*;*Foxj1-CreER^T2^* mice crossed with *Scgb1a1-CreER^TM^*;*R26^RtdTomato^* mice (Fig. 4d). Of note, in the distal airway, *Il1r1* only marks ciliated cells, not secretory cells^30^. Moreover, ciliated cells are known to lack hallmark features of progenitor cells including self-renewal activity and generation of other lineages, including secretory and AT2 cells, which was also supported by our organoid assay (Extended Data Fig. 7f)^30–32^. Given these pieces of evidence, we examined the expression of Notch ligands (*Jag1*, *Jag2*, *Dll1*, *Dll4*) in ciliated cells of *Il1r1^flox/flox^*;*Scgb1a1-CreER^T2/+^*;*R26^RtdTomato/+^* mice after PBS or bleomycin treatment. Consistently, qPCR analysis revealed significant downregulation of *Jag1/2* expression in ciliated cells after injury, indicating a transient reduction in Notch activity as an initial step in secretory cell-mediated regeneration after injury (Fig. 4e). However, ciliated cells of *Foxj1-CreER^T2^*;*Il1r1^flox/flox^*;*Scgb1a1- CreER^T2/+^*;*R26^RtdTomato/+^* mice failed to downregulate the expression of these ligands after bleomycin treatment, suggesting that IL-1β signalling via *Il1r1* mediates the reduction of *Jag1/2* expression in ciliated cells after injury (Fig. 4e). Significantly, after injury, Notch activity was reduced in secretory cells, which was impaired by *Il1r1* deletion in ciliated cells (Fig. 4f-h). We then checked the lineage behaviour of lineage-labelled cells in these mice at day 28 after injury. There was no discernible alteration in lineage-labelled cells by *Il1r1* deletion in ciliated cells in PBS control lungs (Extended Data Fig.7g-i). However, we found a significant reduction in the frequency of lineage-labelled AT2 cells in the lungs of *Foxj1- CreER^T2^*;*Il1r1^flox/flox^*;*Scgb1a1-CreER^T2/+^*;*R26^RtdTomato/+^* mice compared to the lungs of *Il1r1^flox/flox^*;*Scgb1a1-CreER^T2/+^*;*R26^RtdTomato/+^* mice after bleomycin injury (Fig. 4i,j). Taken together, these data indicate that IL-1β signals build niche environments governing the expression of Notch ligands, Jag1 and Jag2, in ciliated cells, which is essential for the fate transition of secretory cells toward AT2 cells during alveolar regeneration.

### Fosl2-mediated AP-1 transcription factor is required for the fate conversion of secretory cells into AT2 cells

To investigate the epigenetic regulation that mediates the differentiation of secretory cells into AT2 cells, we performed ATAC-seq (Assay for Transposase-Accessible Chromatin with high-throughput sequencing) with secretory and secretory-derived AT2 cells (referred to as sAT2 from here). We generated *Sftpc-dsRed^IRES-DTR^* AT2 reporter mice to monitor SPC-expressing AT2 cells and crossed them with the *Scgb1a1-CreER^TM^*;*R26R^fGFP^* mice to trace cells derived from secretory cells after bleomycin injury (Fig. 5a). As expected, lineage-labelled sAT2 cells (GFP^+^dsRed^+^) were increased compared to control at day 28 post injury (Extended Data Fig. 7j). For ATAC-seq analysis, we isolated lineage-labelled secretory (GFP^+^dsRed^−^) and sAT2 (GFP^+^dsRed^+^) cells (Fig. 5a,b). We identified a total of 54,845 and 52,763 peaks in secretory and sAT2 cells, respectively (Fig. 5c). Spearman correlation coefficients showed the high correlation between biological replicates (Extended Data Fig. 8a). The distribution of peaks from secretory and sAT2 cells were similarly represented, and more than half of them fell into enhancers (intron, exon, and distal intergenic elements) (Extended Data Fig. 8b). While approximately 45,548 peaks were common, 9297 and 7316 peaks were mapped as cell-type specific to secretory and sAT2 cells, respectively (Fig. 5c and Extended Data Fig. 8c). Analysis of Gene Ontology (GO) terms with the genes associated with cell-type specific ATAC-seq peaks revealed the distinct characteristics of secretory or AT2 cells (Extended Data Fig. 8d). Indeed, known-secretory cell markers, such as *Gabrp* and *Cyp2f2*, were associated with secretory cell specific-differential peaks, whereas AT2 cell markers, such as *Sftpc* and *Lyz2*, were associated with sAT2 cell-specific peaks (Extended Data Fig. 8e).

**Figure 5.**
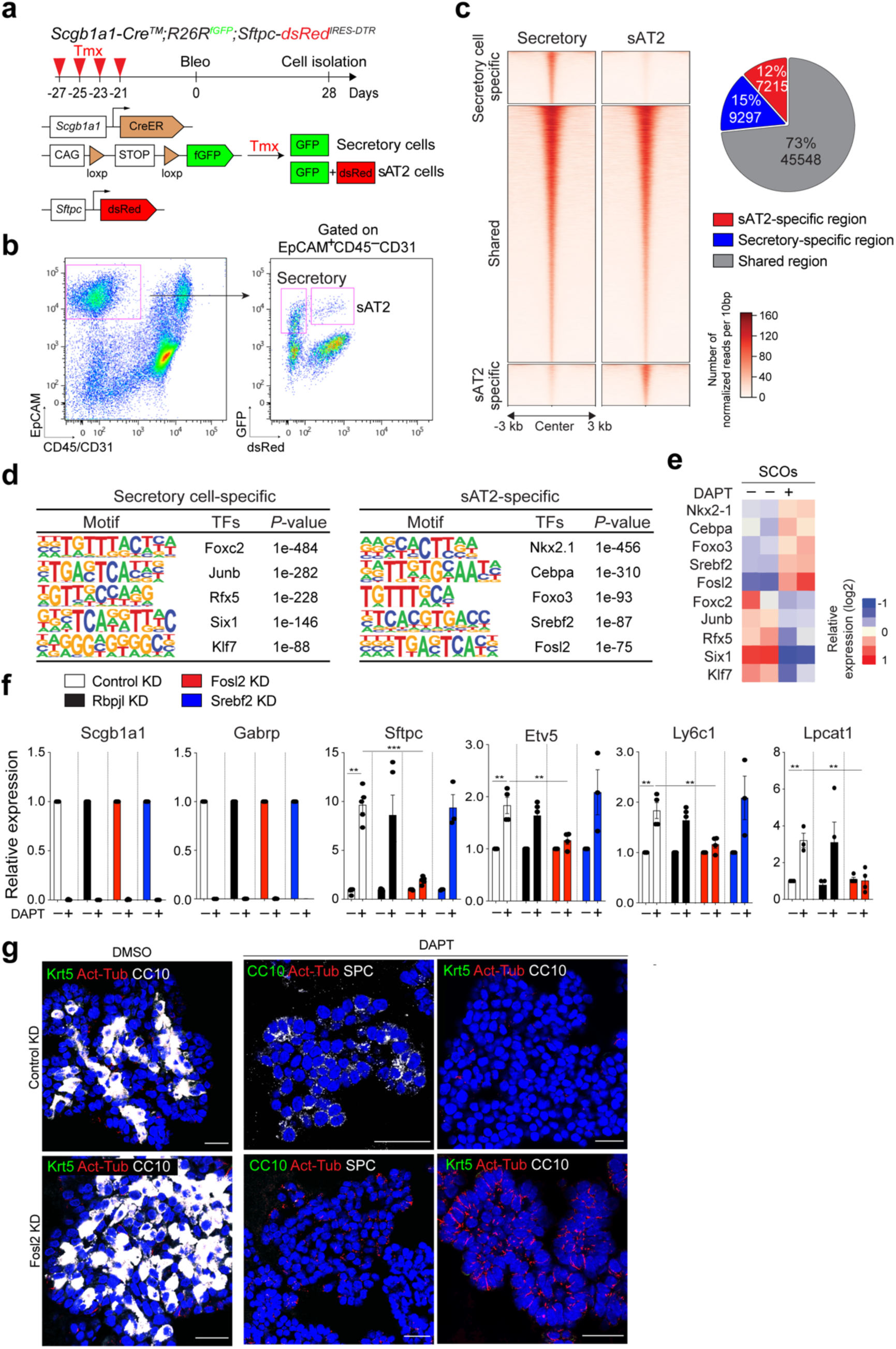
Fosl2/Fra2-mediated AP-1 activity is required for the fate conversion of secretory cells into AT2 cells. **a,** Experimental design for isolation of secretory (GFP^+^dsRed^−^) and secretory-derived AT2 (sAT2, GFP^+^dsRed^+^) cells using *Scgb1a1-CreER^TM/+^*;*R26R^fGFP/+^*;*Sftpc-dsRed^IRES-DTR/+^* mouse post bleomycin injury. Specific time points for tamoxifen injection and isolation of cells are indicated. **b,** Representative flow cytometry analysis for isolation of secretory (GFP^+^dsRed^−^) and sAT2 (GFP^+^dsRed^+^) cells. **c,** A heatmap showing ATAC-seq peaks representing open chromatin regions in secretory cells (left) and sAT2 cells (right). Secretory-specific, shared, and sAT2-specific peaks are shown. Regions of a pie chart presenting the proportion of secretory-specific (blue), sAT2-specific (red), and shared regions (grey). **d,** Top five motif matrices and transcription factors predicted by the HOMER *de novo* motif analysis using secretory-specific and sAT2-specific open regions. **e,** A heatmap showing the expression levels of transcription factors predicted by the motif analysis (Fig. 5d) in secretory cell-derived organoids (SCOs) with or without DAPT treatment (20µM). **f,** qPCR analysis of the markers for secretory (*Scgb1a1* and *Gabrp*) and AT2 cells (*Sftpc*, *Etv5*, *Ly6c1*, and *Lpcat1*) in SCOs with or without DAPT treatment after knockdown of control (white), Rbpjl (black), Fosl2 (red), and Srebf2 (blue). **p<0.01, ***p<0.001. **g,** Representative IF images of secretory cell- derived control KD or *Fosl2* KD organoids treated with DMSO or DAPT (20µM). Single cells dissociated from SCOs at passage 10-12 were cultured in feeder-free condition for 7 days and further maintained with DMSO or DAPT for another 7 days. CC10 (white, left and right panels; green, middle panel), SPC (white, middle panels), Krt5 (green, left and right panels), Act-Tub (red), and DAPI (blue). Scale bars, 50µm.

We sought to identify transcription factors (TFs) that regulate these differential regions of peaks by analysing TF binding motifs. Motif analysis of DNA binding-site showed that sAT2- enriched chromatin contains motifs for key transcriptional factors associated with lung development, including Nkx2-1 and Fosl2^33, 34^ (Fig. 5d). We compared the expression of TFs obtained from motif analysis with RNA-seq data of control and DAPT-treated SCOs (Fig. 2d). The heatmap showed that TFs in sAT2-enriched open chromatin were also highly expressed in DAPT-treated organoids, whereas the expression levels of TFs identified in secretory cell-restricted motif were higher in control organoids (Fig. 5e and Extended Data Fig. 8f). Notably, we found the differential enrichment of AP-1 factor components in control and DAPT-treated organoids; *c-Fos*, *Fosb*, *c-Jun*, and *Junb* in control organoids and *Jund* and *Fosl2/Fra2* in DAPT-treated organoids (Fig. 5e). To interrogate the key TFs that are required for AT2 cell differentiation, we established feeder-free SCOs with KD of five TFs (*Srebf2*, *Fosl2*, *Rbpjl*, *Cebpa*, *Etv5*) selected from motif analysis or a list of differentially expressed genes in RNA-seq analysis. We first validated the KD effect of *Cebpa* and *Etv5*, well-known factors for AT2 cell identity^35, 36^, on the fate switch of secretory to AT2 cells by exposing organoids to DAPT. The upregulation of AT2 cell markers by Notch inhibition was not affected by the interference of *Cebpa* or *Etv5* (Extended Data Fig. 8g,h). The KD of *Srebf2* and *Rbpjl*, which are highly ranked in motif analysis, also did not impair the expression of AT2 cell markers by DAPT treatment (Fig. 5f). However, remarkably, KD of *Fosl2/Fra2* significantly blocked the increase of AT2 cell markers, including *Sftpc* and *Etv5*, by DAPT treatment whereas the downregulation of secretory cell markers such as *Scgb1a1*was not altered (Fig. 5f). Further IF analysis also confirmed impaired induction of SPC^+^ AT2 cells in *Fosl2*-deficient secretory organoids treated with DAPT (Fig. 5g). Interestingly, we found that *Fosl2* KD markedly increased Act-Tub^+^ ciliated cells in DAPT-treated organoids (Fig. 5g). These data suggest that Fosl2 is a critical mediator regulating the fate conversion of secretory cells to AT2 lineage upon Notch inhibition.

### Airway-derived AT2 cells are distinct from alveolar resident AT2 cells in genetic and epigenetic characteristics

While AT2 cells serve as stem cells to maintain the alveolar epithelium during both homeostasis and regeneration, secretory cells are the major source of replenishing alveolar lineages after severe alveolar damage such as bleomycin injury^5, 6^. However, it remains unexplored whether sAT2 cells generated after injury-repair are genetically or functionally identical to resident AT2 cells (hereafter referred to as rAT2 cells) in alveoli. To address this question, we further analysed the AT2 cell population in our scRNA-seq (Fig. 3j) of *Scgb1a1- CreER^TM/+^*;*Red2-Notch^N1ICD/+^* mice challenged by bleomycin injury and found that secretory lineage-labelled AT2 (sAT2, YFP^+^) and non-lineage-labelled AT2 (rAT2, RFP^−^YFP^−^) cells were indeed segregated (Fig. 6a). Notably, sAT2 cells still retained higher expression of many secretory cell markers such as *Scgb1a1*, *Cyp2f2*, and *Sox2* while the expression of AT2 markers such as *Sftpc* and *Etv5* was comparable to that seen in rAT2 cells (Fig. 6b,c). We then examined the epigenetic signatures of sAT2 and rAT2 cells via ATAC-seq on isolated GFP^+^dsRed^+^ (sAT2) and GFP^−^dsRed^+^(rAT2) cells from *Scgb1a1-CreER^TM/+^*;*R26R^fGFP/+^*;*Sftpc-dsRed^IRES- DTR/+^* mice treated with bleomycin injury at day 28 (Extended Data Fig. 9a,b). We identified 52,497 shared open regions from sAT2 and rAT2 cells and 429 differential open regions of sAT2 cells (Extended Data Fig. 9c). Most sAT2-specific open regions were located far away from the transcription start sites (TSS), suggesting that the differential peaks co-localized with contained distal enhancer regions (Extended Data Fig. 9d). Consistent with genetic signatures, the regulatory regions surrounding secretory cell marker loci, including *Scgb1a1* and *Sox2*, showed accessible chromatin signatures in sAT2 cells, much like in secretory cells (Fig. 6d). GO term analysis with the sAT2-specific peaks also supported the enriched secretory-like signature shown in Extended Data Fig. 8d (e.g. tube morphogenesis-related pathway) in sAT2 cells (Extended Data Fig. 9e). Importantly, we found the distinctive feature of an apoptosis-related signature in sAT2 cells (Extended Data Fig. 9e). Genes involved in the negative regulation of apoptotic processes (GO:0043066) were also more highly expressed in sAT2 than rAT2 cells in our scRNA-seq analysis (Extended Data Fig. 9f,g). We confirmed that the *Nr4a2* locus, one of the most critical anti-apoptotic regulators in the lung, is open in sAT2 cells, but not in rAT2 cells^37, 38^ (Extended Data Fig. 9h). Furthermore, sAT2 cells showed higher expression of *Slc7a11*, which is a master regulator for redox homeostasis and essential for survival in response to cellular stress such as cystine deficiency, which can lead to cell death, referred to as ferroptosis^39, 40^ (Fig. 6b).

**Figure 6.**
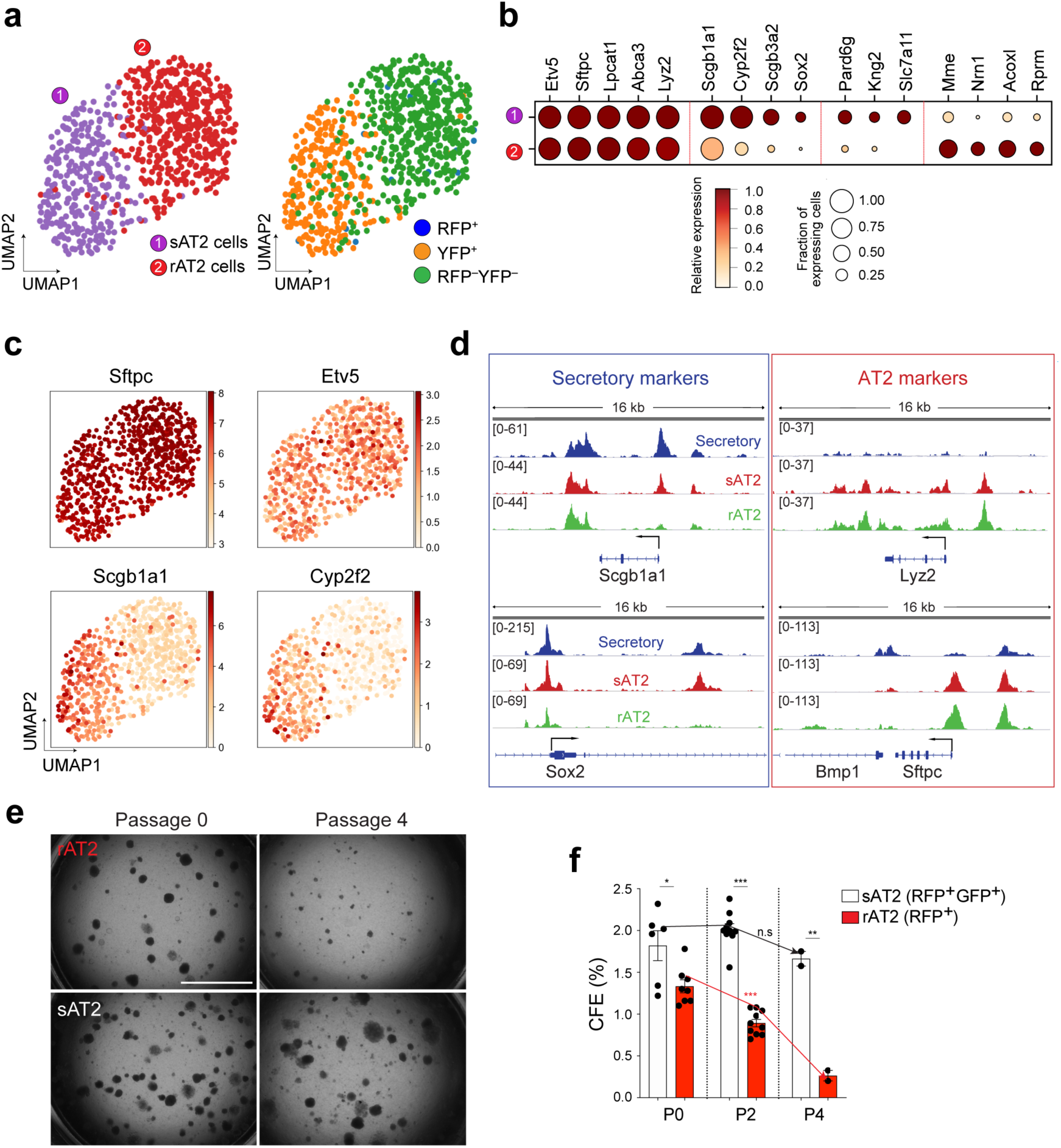
Distinctive features of secretory-derived AT2 cells. **a,** UMAP visualisation of two distinctive AT2 cell clusters sAT2 (secretory cell-derived AT2, YFP^+^SPC^+^ cluster); rAT2 (resident AT2, YFP^−^RFP^−^ non-lineage labelled SPC^+^ cluster) from scRNA-seq analysis of *Scgb1a1-CreER^TM/+^*;*Red2-Notch^N1ICD/+^* mice in Fig. 3j. **b,** Gene expression of key markers in sAT2 and rAT2 cell clusters. **c,** UMAP visualisation of the log- transformed (log10(TPM+1)), normalised expression of selected marker genes (*Sftpc* and *Etv5* for AT2 cells. *Scgb1a1* and *Cyp2f2* for secretory cells) in distinctive clusters shown in (**a**). **d,** Signal track images showing open regions of markers for secretory (Scgb1a1 and Sox2) and AT2 cells (Lyz2 and Sftpc) mapped in secretory (blue), sAT2 (red), and rAT2 cells (green) from the result of Fig. 5c. **e,** Representative bright-field images of organoids derived from rAT2 (top) or sAT2 (bottom) cells: Experiment scheme for labelling with tamoxifen is same as Fig. 5a. *Scgb1a1-CreER^TM/+^*;*R26R^fGFP/+^*;*Sftpc-dsRed^IRES-DTR/+^* mice was given four doses of tamoxifen followed by bleomycin injury. Isolated GFP^+^dsRed^+^ (sAT2) and GFP^−^dsRed^+^ (rAT2) cells at 2 months post bleomycin injury were co-cultured with stromal cells with 1:5 ratio. Images show the organoids at passage 0 and 4. Scale bar, 2,000µm. **f,** Statistical quantification of colony forming efficiency (CFE) of indicated organoids. Each individual dot represents individual biological replicate and data are presented as mean and s.e.m. *p<0.05, **p<0.01, ***p<0.001.

Given these findings, to interrogate the functional differences between sAT2 and rAT2 cells, we established organoid co-cultures of GFP^+^dsRed^+^ (sAT2) and GFP^−^dsRed^+^ (rAT2) cells isolated from *Scgb1a1-CreER^M2/+^*;*R26R^fGFP/+^*;*Sftpc-dsRed^IRES-DTR/+^* mice at 2 months post bleomycin injury. Consistent with previous research^41^, rAT2 cells lost colony forming efficiency (CFE) after serial passaging (Fig. 6e,f). However, sAT2 cells formed stable organoids without loss of CFE over multiple passages (Fig. 6e,f). Moreover, sAT2-derived organoids revealed a greater ratio of AT1 versus AT2 cells compared to that of rAT2-derived organoids, suggesting enhanced differentiation capacity into AT1 lineages (Extended Data Fig. 9i-k). We were also able to detect sAT2 cells (GFP^+^dsRed^+^) in the lungs of *Scgb1a1- CreER^TM/+^*;*R26R^fGFP/+^*;*Sftpc-dsRed* mice for least at 3 months post bleomycin injury (Extended Data Fig. 9l,m). Taken together, these data indicate that AT2 cells derived from secretory cells during alveolar regeneration remain functionally and epigenetically distinct compared to alveolar resident AT2 cells.

### Conserved differentiation potential of human secretory cells marked by KDR/FLK-1

The potential of airway cell compartments to contribute to alveolar regeneration in the human lung has remained unknown. We therefore asked whether secretory cells in the human distal lungs have the potential to convert into AT2 cells, governed by Notch signalling, similar to that observed in the mouse lung. By combining analysis of our scRNA-seq and ATAC-seq analysis, we identified a surface marker *Kdr/Flk-1* that is specifically expressed in mouse secretory cells (Extended Data Fig.10a,b), which is also supported by previous studies^42, 43^. Flow cytometry analysis confirmed that Kdr expression specifically marks secretory cells in the mouse lung (Extended Data Fig.10c-e). We then analysed recently-published scRNA-seq data^44, 45^ and found that *KDR* is also highly expressed in secretory cells in adult human lung tissue (Extended Data Fig. 10f,g). We further confirmed the expression of KDR in secretory cells of distal human lung tissue by IF staining (Fig. 7a). In combination with a human AT2 cell-specific surface marker HTII-280, we were able to sort KDR^+^HTII-280^−^ cells by flow cytometry analysis (Fig. 7b). Further analysis of cytospin staining on freshly isolated KDR^−^HTII-280^+^ and KDR^+^HTII-280^−^ populations revealed that CC10^+^ secretory cells were specifically enriched in the KDR^+^ population (Fig. 7c,d). We also confirmed that KDR^+^HTII-280^−^ cells showed higher expression of secretory cell markers, such as *SCGB1A1* and *SCGB3A2*, and lower expression of AT2 cell markers, such as *SFTPC*, compared to KDR^−^HTII-280^+^ AT2 cells (Fig. 7e). Furthermore, KDR^+^HTII-280^−^ secretory cells could form organoids consisting of mainly CC10^+^ secretory cells and some Act-Tub^+^ ciliated cells, with a few KRT5^+^ basal cells in our culture conditions (Fig. 7f, **See Method**).

**Figure 7.**
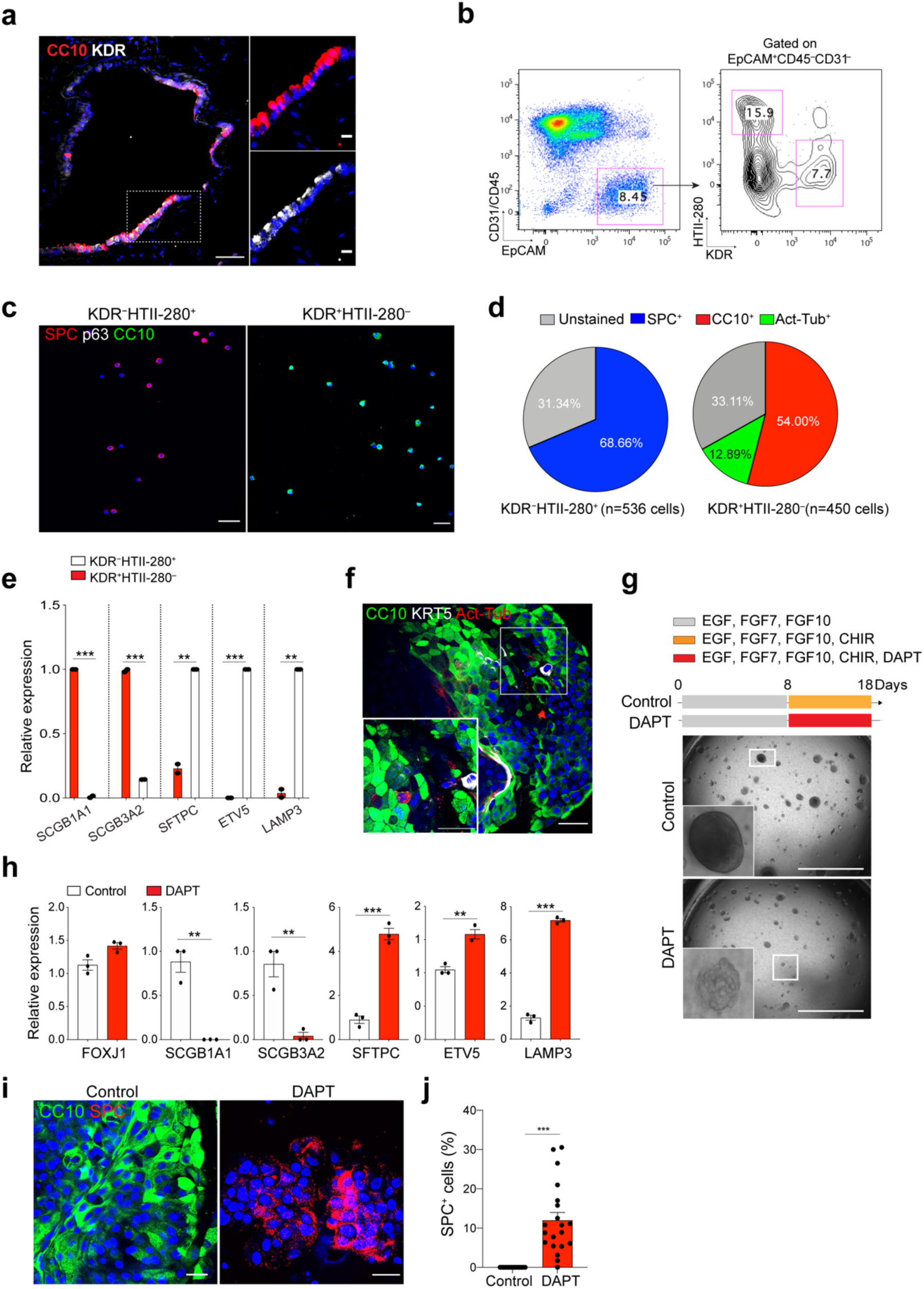
Differentiation plasticity of secretory cells into AT2 cells by inhibition of Notch signalling in human lungs. **a,** IF images showing the expression of KDR in secretory cells in the lung from normal donors. CC10 (red), KDR (white), and DAPI (blue). White boxed insets show high-power view. Scale bar, 50µm and 10µm (inset). **b,** Representative flow cytometry analysis for isolation of KDR^+^HTII-280^−^ or KDR^−^HTII-280^+^ cells from human lung parenchymal tissues. **c,** Representative IF images of cytospin staining from freshly sorted KDR^−^HTII-280^+^ or KDR^+^HTII-280^−^ population. SPC (for AT2 cells, red), p63 (for basal cells, white), CC10 (for secretory cells, green) and DAPI (blue). Scale bars, 100µm. **d,** Quantification of SPC^+^, CC10^+^, and Act-Tub^+^ cells revealed in cytospin staining from KDR^−^HTII-280^+^ or KDR^+^HTII-280^−^ cells in Fig. 7c. **e,** qPCR analysis of the markers for secretory (*SCGB1A1* and *SCGB3A2*) and AT2 (*SFTPC*, *ETV5*, and *LAMP3*) cells in freshly isolated KDR^+^HTII-280^−^ (red bars) or KDR^−^HTII-280^+^ (white bars) cells. **p<0.01, ***p<0.001. **f,** Representative IF images of KDR^+^ cell-derived organoids at first passage. KDR^+^HTII-280^−^ cells were cultured in the basic medium supplemented with EGF, NOG, FGF7, and FGF10 for 3 weeks. CC10 (green), Act- Tub (red), KRT5 (white), and DAPI (blue). Scale bar, 50µm. **g,** Representative bright-field images of organoids derived from KDR^+^HTII-280^−^ cells with or without DAPT treatment (20µM) at passage 1. Isolated KDR^+^HTII-280^−^ cells were cultured with airway cell culture medium (See **Methods**) supplemented with EGF (50ng/ml), FGF7 (100ng/ml), and FGF10 (100ng/ml) for 8 days, followed by inclusion of CHIR99021 (2µM) for additional 10 days with or without DAPT (20µM). Scale bar, 2,000µm. **h**, qPCR analysis of the markers for secretory cells (*SCGB1A1* and *SCGB3A2*) and AT2 cells (*SFTPC*, *ETV5*, and *LAMP3*) in organoids derived KDR^+^HTII-280^−^ cells without (white bars) or with DAPT treatment (red bars). Each individual dot represents individual biological replicate and data are presented as mean ± s.e.m. **p<0.01, ***p<0.001. **i**, Representative IF images of KDR^+^ cell-derived organoids treated with DMSO or DAPT (20µM). CC10 (green), SPC (red), and DAPI (blue). Scale bar, 50µm. **j**, Quantification of the frequency of SPC^+^ cells in the control or DAPT treated organoids. Each individual dot represents one organoid and data are presented as mean ± s.e.m (n=3 independent biological replicates). ***p<0.001.

Given that human secretory cells also showed higher Notch activity (e.g. *HES1* expression) similar to mouse tissue (Extended Data Fig. 10f), we inhibited Notch activity by treating organoids derived from KDR^+^HTII-280^−^ cells with DAPT to investigate the differentiation potential of human secretory cells into AT2 cells. Notably, similar to mouse secretory cells, Notch inhibition induced the morphological changes of human secretory cell derived-organoids with significant upregulation of AT2 marker expression, such as *SFTPC*, whereas the expression of secretory cell markers, including *SCGB1A1* was significantly reduced (Fig. 7g,h). IF staining also confirmed that Notch inhibition drives the generation of SPC^+^ AT2 cells at the expense of SCGB1A1/CC10^+^ secretory cells in DAPT-treated organoids (Fig. 7i,j). We barely observed any significant alterations in other lineages such as Act-Tub^+^ ciliated cells and KRT5^+^basal cells (Extended Data Fig. 10h-j). Overall, our data suggest that KDR^+^ secretory cells share a conserved key molecular component of Notch signalling in governing alveolar regeneration of human and mouse lungs.

## Discussion

Spatiotemporal dynamics of stem and progenitor cells ensure the rapid and efficient process of tissue repair for the reconstruction of epithelial integrity and function. In response to severe lung injury, secretory cells localised in the distal airway mobilise and differentiate into AT2 cells to compensate for the loss of alveolar epithelium and restore alveolar function. However, the cellular events and regulatory networks directing the differentiation plasticity of secretory cells, and the identity of the functional niches during this process, have been largely elusive. Particularly, how secretory cells are activated and acquire differentiation plasticity is unknown. Here, our data reveal two sequential stages of secretory cell fate conversion into AT2 cells during alveolar regeneration. Loss of Notch activity mediated by IL-1β signalling in ciliated cells reprograms the secretory cells to lose their identity upon injury. Fosl2/Fra2-mediated AP- 1 transcription factor then drives the fate conversion of secretory cells into AT2 cells. Furthermore, using scRNA-seq and ATAC-seq analysis, we identified that secretory-derived AT2 cells post injury retain the airway lineage identity in both a genetic and epigenetic manner, and have greater functional capacity of long-term maintenance *in vitro* compared to resident AT2 cells. By identifying a surface marker KDR/FLK-1 of human secretory cells, we propose that functionally equivalent human secretory cells are capable of contributing to alveolar lineages post injury via Notch signalling.

In this study, we uncovered the role of Notch signalling as a key regulator in reprogramming secretory cells to acquire differentiation plasticity post alveolar injury. Inhibition of Notch signalling resulted in the loss of secretory cell identity and simultaneously enhanced the fate conversion to AT2 cells upon injury. In contrast, constitutive activation of Notch signalling substantially blocked AT2 cell differentiation from secretory cells post injury, indicating that regulation of Notch activity is critical for regeneration in the context of the lung. Instead, cells were arrested at the intermediate cell state expressing high levels of *Cdkn1c/p57* expression, which is likely to link the fate transition between secretory and AT2 cells. These results reveal that the downregulation of Notch activity is indispensable for AT2 cell differentiation from secretory cells during regeneration. Notably, the intermediate population is specifically marked by the expression of *Fstl1* that has been known to antagonize BMP signalling^46^. Given the functional role of BMP signalling in facilitating proximal epithelium with a concurrent reduction in the distal epithelium during lung development^46, 47^, impaired alveolar regeneration in constitutive Notch activity could be connected to compromised proximal-distal patterning, which may be due to inhibition of BMP signalling. It would be interesting to further define the presence of *Fstl1*-like populations in lung development, and whether alveolar regeneration reflects the developmental stages.

How is the activity of the Notch pathway in secretory cells controlled? Given the communication characteristics of Notch signalling via direct contact, we identified ciliated cells as functional niches to induce Notch signalling in secretory cells by expressing Notch ligands, including Jag1, which is also consistent with a previous study in the developing lung^29^. An increasing body of evidence is revealing that inflammation induced by tissue injury modulates repair and tissue regeneration^48–50^. Indeed, our recent study showed that acute inflammatory niches driven by IL-1β are essential for alveolar regeneration^30^. Of note, we found that *Il1r1* is highly expressed in ciliated cells, and the expression of Notch ligands on ciliated cells is regulated by IL-1β signalling. Specific deletion of *Il1r1* in ciliated cells significantly impaired the contribution of secretory cells in alveolar reconstruction. This is suggestive that an IL-1β-mediated inflammatory niche coordinates alveolar regeneration by regulating the behaviour and cell fate transition of distal secretory cells, in addition to subsets of *Il1r1^+^* AT2 cells post injury^30^.

By combining ATAC-seq and RNA-seq analysis, we identified Fra2/Fosl2, which together comprise the heterodimeric AP-1 transcription factor, as an essential driver for the fate conversion of secretory cells into AT2 lineages. Notably, we found that the components of the AP-1 family were differentially expressed in control and DAPT-treated organoids derived from secretory cells. While the expression of *c-Jun*, *c-Fos*, *Junb*, and *Fosb* was downregulated by DAPT treatment, the expression of *Fosl2* and *Jund* were specifically upregulated. Further, the Harmonizome database also shows that Fosl2 is a functional target of Rbpj, a key mediator of Notch signalling, and several regulatory regions of AT2 markers, including *Sftpc* and *Etv5*, are directly occupied by *Fosl2*^51^. Importantly, *Fosl2*-deficient secretory cells failed to acquire AT2 cell differentiation in the presence of DAPT, however decreased secretory cells were not recovered. Further, *Fosl2* KD didn’t influence the cell fate of secretory cells without Notch inhibition. These data suggest that Notch inhibition coordinates the cell fate decisions of secretory cells into AT2 lineages in two stages: 1) loss of secretory cell identity and 2) acquisition of transcriptional programmes for AT2 cell differentiation (Extended Data Fig. 10k). Our data revealed that Fosl2 is required for AT2 cell conversion from secretory cells, but dispensable for the loss of secretory cell identity. Other candidates, such as *Foxc2* and *Six1*, that occupy the regulatory region of secretory cell-specific genes and are substantially downregulated by Notch inhibition (Fig.5d), could be tested for potential roles in regulating the maintenance of secretory cell identity.

Notch inhibition has been reported to promote the differentiation of secretory cells into ciliated cells in lung development and homeostatic adult lungs, although there are some differences in the incidence of ciliated cell differentiation, potentially due to different contexts or model systems^25–27^. In this study, we demonstrated that, during alveolar regeneration after injury, Notch inhibition facilitated the fate conversion of secretory cells into AT2 cells via *IL-1β- Notch-Fosl2* axis. Despite increased ciliated cell differentiation of *dnMAML*-expressing secretory cells prior to injury, persistent Notch inhibition enhanced AT2 cell differentiation via *Fosl2* during injury repair. How does *Fosl2* regulate this process? Interestingly, withdrawals of WNT signalling-inducing factors, including Wnt3a and R-spondin-1, in our organoid cultures, showed a dramatic increase in Act-Tub^+^ ciliated cells at the expense of secretory cells in DAPT-treated organoids (Extended Data Fig.10l). AT2 cells were barely observed in these organoids (Extended Data Fig.10l). This result suggests that Notch and WNT signalling likely function cooperatively in secretory cells differentiating to AT2 cells. Indeed, previous studies showed increased expression of Wnt ligands in mesenchymal cells during alveolar regeneration after injury^19, 52^. Further analysis of scRNA-seq from our previous study^30^ revealed that Wnt ligands, such as *Wnt5a* and *Wnt7b*, were highly upregulated at day 14 and returned back to homeostatic levels at day 28 post bleomycin injury (Extended Data Fig.10m). Notably, however, *Fosl2*-KD secretory cells failed to generate AT2 cells even in the presence of Wnt activity (Fig. 5g). Given these results, it is likely that increased Wnt ligands from mesenchymal cells, in parallel with Notch-mediated Fosl2 activity, may cooperatively confer differentiation potential of secretory cells into AT2 lineages, via independent regulatory axes, during alveolar injury repair. While Fosl2 was identified as a key factor in the axis of Notch signalling, further identification of interacting partners regulated by Wnt signalling will uncover the molecular networks governing the fate decision of secretory cells into ciliated or AT2 cells in homeostasis and regeneration.

Increasing evidence reveals the plasticity of epithelial stem cells under regenerative conditions. In the distal lungs, two stem cell compartments, secretory cells and AT2 cells, maintain the airway and alveolar epithelium, respectively, in homeostasis and regeneration. However, upon severe damage, secretory cells acquire differentiation plasticity to generate alveolar lineages. Interestingly, we found that sAT2 cells display distinct features compared to resident AT2 (rAT2) cells, even after injury resolution. sAT2 cells still retained some airway lineage identity based on the higher expression of secretory cell markers such as *Scgb1a1* and *Sox2*, whereas most genetic and epigenetic signatures of genes, including canonical AT2 cell markers such as *Sftpc* and *Etv5*, are shared with rAT2 cells. Notably, sAT2 cells are capable of long-term self-renewal compared to rAT2 cells in *in vitro* organoids, which may be attributed to the enriched signatures of anti-apoptotic functions such as *Slc7a11* and *Nr4a2* in sAT2 cells compared to rAT2 cells. sAT2 cells may have a higher tolerance for severe alveolar insults than rAT2 cells, enabling them to replenish damaged alveolar epithelium more efficiently. Notably, in chronic lung diseases such as pulmonary fibrosis and lung cancer, the destruction of alveolar structure with bronchiolization is a common feature^53, 54^. Moreover, AT2 cells expressing airway- restricted genes such as Sox2 were also seen in fibrosis patients^55^. Given our findings, it would be interesting to explore further whether the genetic and epigenetic memory of sAT2 cells contributes to this pathologic phenotype after injury resolution.

The loss of alveolar integrity is a life-threatening and key pathologic feature of various chronic lung diseases. Recent studies suggested the regenerative potential of human HTII-280^+^ AT2 cells using an *in vitro* organoid model, yet their capacity to generate new alveolar epithelium seems limited^22, 23^. In particular, the functional capacity of human airway secretory cells to contribute to alveolar lineages has never been explored, largely due to the lack of information for isolation and culture of secretory cells *in vitro*. Here we provide the first evidence of the potential progenitor capacities of secretory cells in self-renewal and generation of AT2 cells using human cells, by identifying KDR/FLK-1 as the surface marker of secretory cells and establishing a feeder-free organoid culture. We furthermore demonstrated that KDR^+^ secretory cells in human lungs have the differentiation plasticity to give rise to AT2 cells, which is also regulated by Notch signalling, as we demonstrated in mouse lungs. Our findings provide a solid foundation for further studies of the potential role of these cells in chronic lung diseases.

In summary, our results identify the molecular and cellular mechanisms of the cross-compartment contribution of airway secretory cells to the regeneration of the alveolar epithelium after lung injury. The presence of conserved markers and ease of cultivation of both mouse and human secretory cells provide a unique opportunity for mechanistic studies to shed light on human lung progenitor cell biology and assist in developing new treatments for acute and chronic lung diseases. Our study also provides clues for the potential therapeutic targets of Notch signalling in lung diseases caused by defective and dysregulated alveolar regeneration.

## Acknowledgement

We would like to thank Emma Rawlins (Gurdon Institute, University of Cambridge, UK) for sharing *Scgb1a1-CreER^TM^*, *Foxj1-CreER^T2^*, and *Rosa26R-CAG-fGFP* mouse line; Irina Pshenichnaya (Histology), Maike Paramor (NGS library), Peter Humphreys (Imaging), Andy Riddell (Flow cytometry), Simon McCallum (Flow cytometry, Cambridge NIHR BRC Cell Phenotyping Hub), Katarzyna Kania (single cell sequencing at Cancer Research UK), William Mansfield (Transgenic mice) and Cambridge Stem Cell Institute core facilities for technical assistance; Jong-Eun Park (Welcome Sanger Institute, UK) and Seungmin Han (Gurdon institute, UK) for helpful discussion on the scRNA-seq analysis; Yujin Ahn (DGIST, Korea) for technical assistance for shRNA experiment. All Lee Lab members for helpful discussion. This work was supported by Wellcome and the Royal Society (107633/Z/15/Z) and European Research Council Starting Grant (679411). J.K. is supported by R01GM112722 from the National Institute of General Medical Sciences and Preterm Birth Research Grant from the Burroughs Wellcome Fund. S.M.J is funded by the Medical Research Council UKRMP2, and the Cancer Research UK Programme grant and Lung Cancer Centre of Excellence. B.D.S acknowledges funding from the Royal Society E.P. Abraham Research Professorship (RP/R1/180165) and Wellcome Trust (098357/Z/12/Z).

## Author contribution

J.C. Y.J.J, J.K., and J.-H.L. designed the experiments, interpreted the data, and wrote the manuscript; J.C. performed most experiments and data analysis; Y.J.J. performed and analysed bulk RNA-seq and ATAC-seq data; C.D. designed and performed lineage-tracing analysis of *Red2-Notch^N1ICD^* mice, provided valuable comments on the manuscript; E. I. generated the *Sftpc-IRES-DTR-P2A-dsRed* targeting vector; K.V.E. performed the isolation of human lung tissue; J.-H.L. generated *Sftpc-IRES-DTR-P2A-dsRed* mouse line; B.-K.K. helped with the generation of the *Sftpc-IRES-DTR-P2A-dsRed* targeting vector and shared the *Red2-Notch^N1ICD^* mice line; B.D.S helped with the study of *Red2- Notch^N1ICD^* lineage-tracing analysis; H.H. and S.M.J provided human lung tissue samples.

## Data and materials availability

All open access-consented sequencing data (i.e. bulk RNA-seq, scRNA-seq, and ATAC-seq) will be deposited upon acceptance of the manuscript.

## Methods

### Animals

*Scgb1a1-CreER^TM^*^13^, *Sftpc-CreER^T2^* ^5^, *Foxj1-CreER^T2^*^31^, *Red2-Notch^N1ICD^* (Yum et al., under revision), *Rosa26R-CAG-fGFP*^13^, *Rosa26R-lox-stop-lox-tdTomato*^56^, and *Il1r1^flox/flox^*^57^ mice have been described and are available from Jackson Laboratory. Il1r1-P2A- eGFP-IRES-CreER^T2^ (*Il1r1-CreER^T2^*)^30^. To monitor SPC-expressing AT2 cells, our lab generated Sftpc-IRES-DTR-P2A-dsRed (*Sftpc-dsRed^IRES-DTR^*) reporter mouse where IRES-DTR-P2A-dsRed construct is inserted into 3’-UTR region of endogenous Sftpc gene. Mice for the lineage tracing and injury experiments were on a C57BL/6 background and 6-10 week old mice were used for most of the experiments described in this study. Experiments were approved by local ethical review committees and conducted according to UK Home Office project license PC7F8AE82. Mice were bred and maintained under specific-pathogen-free conditions at the Cambridge Stem Cell Institute and Gurdon Institute of University of Cambridge.

### Tamoxifen

Tamoxifen (Sigma) was dissolved in Mazola corn oil (Sigma) in a 20mg/ml stock solution. 0.2mg/g body weight tamoxifen was given via intraperitoneal (IP) injection. The numbers and date of treatment are indicated in the individual figures of each experimental scheme.

### Bleomycin administration

6-10 week-old mice were anesthetised via inhalation of isoflurane for approximately 3 mins. The mice were positioned on the intratracheal intubation stand, and 1.25U/kg of bleomycin, or PBS control, were delivered intratracheally by a catheter (22G). During the procedure anaesthesia was maintained by isoflurane and oxygen delivery.

### Mouse lung tissue dissociation and flow cytometry

Lung tissues were dissociated with a collagenase/dispase solution as previously described^30^. Briefly, after lungs were cleared by perfusion with cold PBS through the right ventricle, 2ml of dispase (BD Biosciences, 50Uml) was instilled into the lungs through the trachea until the lungs inflated. Each lobe was dissected and minced into small pieces in a conical tube containing 3 ml of PBS, 60µl of collagenase/dispase (Roche), and 7.5µl of 1% DNase I (Sigma) followed by rotating incubation for 45 min at 37°C. The cells were then filtered sequentially through 100- and 40-µm strainers and centrifuged at 1500 rpm for 5 min at 4°C. The cell pellet was resuspended in 1ml of RBC lysis buffer (ACK buffer, 0.15M NH4Cl, 10mM KHCO3, 0.1mM EDTA) and lysed for 2 min at room temperature. 6 ml basic F12 media (GIBCO) was added and 500µl of FBS (Hyclone) was slowly added in the bottom of tube. Cells were centrifuged at 1500 rpm for 5 min at 4°C. The cell pellet was resuspended in PF10 buffer (PBS with 10% FBS) for further staining. The antibodies used were as follows: CD45 (30-F11)-APC or -APC-Cy7 (BD Biosciences), CD31 (MEC13.3)-APC (BD Biosciences), EpCAM (G8.8)-PE-Cy7 or FITC (BioLegend), and CD24 (M1/69)-APC (eBioscience), MHC-II (I-A/I-E, M5)-FITC or -APC-Cy7 (eBioscience), and CD309/Kdr/Flk-1 (7D4-6)-APC (BioLegend). 4’, 6-diamidino-2-phenylindole (DAPI) (Sigma) was used to eliminate dead cells. Data were acquired on LSRII analyser (BD Bioscience) and then analysed with FlowJo software (Tree Star). MOFLO system (Beckman Coulter) was used for the sorting at Cambridge NIHR BRC Cell Phenotyping Hub.

### Human lung tissue dissociation and flow cytometry

Human lung tissues were cleared by perfusion with cold PBS and cut into small (1 mm) pieces. Small pieces of lung tissues (2 cm x 2 cm) were moved into a conical tube containing 10 ml of digestion buffers (2ml of dispase II (Sigma), 100 µl of collagenase/dispase (Roche), 100 µl of 1% DNase I (Sigma), and 7.8 ml of PBS), followed by rotating incubation for 1 hr at 37 °C. The cells were then filtered through 100 µm strainers and centrifuged at 350 g for 5 min at 4°C. The cell pellet was resuspended in 5 ml of RBC lysis buffer (ACK buffer, 0.15 M NH4Cl, 10 mM KHCO3, 0.1 mM EDTA) and lysed for 5 min at room temperature. 10 ml basic F12 media (GIBCO) was added and 500 µl of FBS (Hyclone) was slowly added in the bottom of tube. Cells were centrifuged at 350g for 5 min at 4°C. The cell pellet was resuspended in PF10 buffer (PBS with 10% FBS) for further staining. The antibodies used were as follows: CD45 (2D1)-APC (BioLegend), CD31 (VM59)- APC (BioLegend), EpCAM (9C4)- FITC (BioLegend), HTII-280-mouse IgM (Terrace Biotech), Purified CD309/KDR (A16085H), anti-mouse IgG1 (RMG1-1)-APC-Cy7 (BioLegend), and anti-mouse IgM (Il/41)-PE (eBioscience). 4’, 6-diamidino-2-phenylindole (DAPI) (Sigma) was used to eliminate dead cells. MOFLO system (Beckman Coulter) or Flexible BD Influx^TM^ cell sorter were used for the sorting at Cambridge NIHR BRC Cell Phenotyping Hub and data were analysed with FlowJo software (Tree Star).

### *In vitro* feeder-free organoid culture and passages

Freshly sorted lineage-labelled cells from *Scgb1a1-CreER^TM^*;*R26R^tdTomato^* or *Sftpc-CreER^T2^*;*R26R^tdTomato^* mice were resuspended in basic medium (AdDMEM/F12 (Invitrogen) supplemented with B27 (Invitrogen), 1mM *N*- Acetylcysteine (Sigma), 10 mM Nicotinamide (Sigma)). Then, cells were mixed with growth factor-reduced Matrigel (BD Biosciences) at a ratio of 1:1. A 100 µl mixture was placed in a 24-well Transwell insert with a 0.4 µm pore (Corning). Approximately 10×10^3^ epithelial cells were seeded in each insert. After Growth Factor Reduced Matrigel (GFR) formed a gel, 500 µl of culture medium (basic medium supplemented with the growth factors: 50 ng/ml murine EGF (Life Technology), 100 ng/ml human FGF7/KGF (Peprotech), 100 ng/ml human FGF10 (Peprotech), 50% WNT3A-conditioned media (provided by Tissue Core Facility of Cambridge Stem Cell Institute), 10% RSPO1-conditioned media (provided by Tissue Core Facility of Cambridge Stem Cell Institute), 100 ng/ml Noggin (Peprotech)) was placed in the lower chamber. Medium was changed every other day and ROCK inhibitor Y27632 (10 µM, Sigma) was added in the medium for the first 2 days of culture. Passage was performed once per 2 weeks at least for 12 months. For passages, organoids were removed from the Matrigel by incubation with dispase (1 mg/ml) for 40 mins at 37 °C, followed by dissociation into single cells using trypLE (Gibco) treatment for 5 min at 37 °C. 5∼10×10^3^ cells were transferred to fresh Matrigel in 24-well Transwell insert. For organoid culture from a single cell with limiting dilution, FACS-sorted cells were plated into 48-well plates (Corning) with one cell per well. Every dilution was replicated in 48-well plates (Corning) for two independent experiments. Single cells imbedded in Matrigel were monitored at microscope with RFP channels to check the expression of Tomato expression.

For human KDR^+^ cell culture in organoids, approximately 10×10^3^ epithelial cells were resuspended in a 20 µl of 100% GFR-Matrigel and seeded in 48 well plates, followed by 30min incubation at 37 °C for solidification. Then, 250µl of airway cell culture medium (basic medium supplemented with the growth factors: murine EGF (50 ng/ml, Life Technology), human FGF7/KGF (100 ng/ml, Peprotech), human FGF10 (100 ng/ml, Peprotech), Noggin (100 ng/ml, Peprotech), A83-01 (1 µM, Tocris), SB202190 (1 µM, Tocris) was added to each well. To avoid the growth of fungal and bacterial infection, 250 ng/mL Amphotericin B and 50µg/mL gentamicin were added to culture medium. For culture with DATP treatment in Fig.7g, organoids cultured for 8 days with airway cell culture medium were followed by inclusion of CHIR99021 (2 µM, Tocris) for additional 10 days with or without DAPT (20 µM). Medium was changed every 3-4 days and ROCK inhibitor Y27632 (10 µM, Sigma) was added in the medium for the first 4 days of culture. Passage was performed once per 3 weeks.

### Knockdown construct and retroviral infection to organoids

For sequence-specific knockdown of candidate genes, target sequences were cloned into MSCV-LTRmiR30-PIG (LMP, Open Biosystems) retroviral vectors or pLKO.1-puro lentiviral vector. To generate retroviruses for infection into cells, the HEK293T cells were transfected using a standard calcium phosphate protocol with vectors expressing GFP alone (Control-RV), GFP plus short-hairpin against target genes. Viral supernatants were harvested 2 days later. Secretory-derived organoids of passages between 15 and 20 were prepared after recovery from Matrigel (BD Biosciences) by treatment of dispase (1mg/ml) for 40 min at 37°C, followed by dissociation into single cells using trypLE (Gibco) treatment for 5 min at 37°C. Single cells dissociated from established organoids were infected with the viral supernatants in the presence of polybrene (Sigma, 8μg/ml) by spin infection for 90 min at 2400rpm at 32°C. This procedure was repeated twice every day. Short-hairpin sequences for target genes are as follows: fgeOshEtv5: 5’- TTCTATGAGCTTAAATTCCGC-3’ shCebpa: 5’-TTTGTTTGGCTTTATCTCCGC -3’ shSrebf2: 5’-TATTTGATGTAATCAATGCGC-3’ shRbpjl: 5’-TTCAAAGTTCAACTTCTGCGC-3’ shFosl2: 5’-TATATCTACCCGGAACTTCGC-3’ shRbpj: 5’-GCAGACTCATTGGGCTACATT-3’

### *In vitro* lung organoid co-culture with established stromal cells

Freshly sorted lineage-labelled cells were resuspended in culture medium (3D basic medium (DMEM/F12, Gibco) supplemented with 10% FBS. (Gibco) and ITS (Insulin-Transferrin-Selenium, Corning)), and mixed with cultured lung stromal cells negatively isolated by microbeads of CD326/EpCAM, CD45, and CD31 via MACS (Miltenyi Biotech), followed by resuspension in growth factor-reduced Matrigel (BD Biosciences) at a ratio of 1:5. A 100 µl mixture was placed in a 24-well Transwell insert with a 0.4 µm pore (Corning). Approximately 5×10^3^ epithelial cells were seeded in each insert. 500 µl of culture medium was placed in the lower chamber, and medium was changed every other day. ROCK inhibitor Y27632 (10 µM, Sigma) was added in the medium for the first 2 days of culture. Analysis of colony forming efficiency (CFE) and size of organoids was performed at 14 days after plating if there is no specific description.

### *In vitro* culture of ciliated cells for IL-1β treatment

CD24^high^Tomato^+^ ciliated cells were isolated from *Il1r1-CreER^T2^* mice at day 4 after four doses of tamoxifen treatment. Purified 20,000 cells were embedded in Matrigel with PBS or IL-1β (10 ng/ml) for 24 hrs. Then, RNA was isolated to analyse the expression of *Jag1* or *Jag2*.

### Transplantation of *Scgb1a1*^+^ lineage-labelled cells

Freshly sorted 20,000 cells of CD45^−^EpCAM^+^Tomato^+^MHCII^−^ cells from *Scgb1a1-CreER^TM/+^*;*R26R^tdTomato/+^* or *dnMAML^flox/+^*;*Scgb1a1-CreER^TM/+^*;*R26R^tdTomato/+^* mice were mixed with lung stromal cells isolated from WT mice (20,000 cells) to support epithelial cell survival during engraftment, and were transplanted into WT C57BL/6 mice at day 7 after bleomycin injury (1.25 U/kg). Lungs were analysed at day 14 post injury to determine the differentiation of engrafted cells.

### Quantitative RT-PCR

Total RNA was isolated using a Qiagen RNeasy Mini-plus Kit according the manufacturer’s instructions. Equivalent quantities of total RNA were reverse-transcribed with SuperScript IV cDNA synthesis kit (Life Technology). Diluted cDNA was analysed by real-time PCR (StepOnePlus; Applied Biosystem). Pre-designed probe sets (Thermo Fisher Scientific) were used as follows: human Scgb1a1 (Hs00171092_m1), human Sftpc (Hs00951326_g1), and Human Foxj1 (Hs00230964_m1). Actb expression (Hs01060665_g1) was used to normalise samples using the ΔCt method. Sybr green assays were also used for human or mouse gene expression with SYBR Green Master Mix (2x, Thermo Fisher Scientific). Primer sequences are as follows:

#### Mouse

Gapdh: F-AGGTCGGTGTGAACGGATTTG, R-TGTAGACCATGTAGTTGAGGTCA Scgb1a1: F- ATGAAGATCGCCATCACAATCAC, R-GGATGCCACATAACCAGACTCT Scgb3a2: F-CCACTGCCCTTCTCATCAACC, R-TGTCGTCCAAAGGTACAGGTA Gabrp: F-CAGACCCACGGCTAGTGTTC, R- AGAGGCGGATGAGCCTGTT Cldn10: F-AATCGTCGCCTTCGTAGTC, R-GTTGGCAAAAATAAGTGGCT Cyp2f2: F- GGACCCAAACCTCTCCCAATC, R- CCGTGAACACCGACCCATAC Sftpc: F-TCAGTCTGATAACTTGGTGCTTC, R-GGCTTCCTATCGTAGGCACAA Etv5: F-TCAGTCTGATAACTTGGTGCTTC, R-GGCTTCCTATCGTAGGCACAA Lamp3: F- TCCAAAAGCCAGAGGCTATCT, R- ACTGGGGTTACTGTTTTCATTGT Lpcat1: F-GGCTCCTGTTCGCTGCTTT, R-TTCACAGCTACACGGTGGAAG Abca3: F-CAGCTCACCCTCCTACTCTG, R-ACTGGATCTTCAAGCGAAGCC Jag1: F-CCTCGGGTCAGTTTGAGCTG, R-CCTTGAGGCACACTTTGAAGTA Jag2: F-CAATGACACCACTCCAGATGAG, R-GGCCAAAGAAGTCGTTGCG Dll-1: F- CAGGACCTTCTTTCGCGTATG, R- AAGGGGAATCGGATGGGGTT Dll-4: F- TTCCAGGCAACCTTCTCCGA, R- ACTGCCGCTATTCTTGTCCC Hes1: F-CCAGCCAGTGTCAACACGA, R-AATGCCGGGAGCTATCTTTCT Nrarp: F-TTCAACGTGAACTCGTTCGGG, R-TTGCCGTCGATGACTGACTG

#### Human

Scgb3a2: F- AAGCTGGTAACTATCTTCCTGCT, R- AGGGGCACTTTGTTGATGAGG Lamp3: F- ACTACCCCAGCGACTACAAAA, R- CTAGGGCCGACTGTAACTTCA Etv5: F- TCAGCAAGTCCCTTTTATGGTC, R- GCTCTTCAGAATCGTGAGCCA

### Cytospin, Immunofluorescence, and immunohistochemistry

Mouse lung tissues were routinely perfused, inflated, and fixed with 4% PFA for 2 hrs at 4°C. Cryosections (8-12µm) and paraffin sections (7µm) were used for histology and Immunofluorescence (IF) analysis. Cultured colonies from organoids were fixed with 4% PFA for 2 hrs at room temperature followed by immobilisation with Histogel (Thermo Scientific) for paraffin embedding. For cytospin staining, isolated 2,000 cells were resuspended in 250µl of PBS supplemented with 10% FBS, followed by spinning in pre-wet cytospin funnels at 600rpm for 5min. Sectioned lung tissues or colonies were stained with hematoxylin and eosin (H&E) or immunostained: after antigen retrieval with citric acid (0.01M, pH 6.0), blocking was performed with 5% normal donkey serum in 0.2% Triton-X/PBS at room temperature for 1hr. Primary antibodies were incubated overnight at 4°C at the indicated dilutions: Goat anti-CCSP/CC10 (T-18) (1:200, Santa Cruz Biotechnology Inc., sc-9772), Mouse anti-CCSP/CC10 (E11) (1:200, Santa Cruz Biotechnology Inc., sc-365992), Goat anti-SP-C (1:200, Santa Cruz Biotechnology Inc., sc-7706), Rabbit anti-pro-SP-C (1:300, Millipore, AB3786), Mouse anti-Acetylated Tubulin (6-11B-1) (1:300, Sigma-Aldrich, T7451), Hamster anti-PDPN (1:300, DSHB, 8.1.1), Rabbit anti-KRT5 (1:300, BioLegend, 905501), Mouse anti-P63 (1:200, Abcam, ab735), Rabbit anti- PORCN (1:100, Abcam, ab201793), Rabbit anti-P57/KIP2 (1:100, Abcam, ab75974), Rabbit anti-RFP (1:250, Rockland, 600–401379), Rabbit anti-HOPX (1:100, Santa Cruz Biotechnology Inc., sc-30216), Rabbit anti-HES1 (1:200, D6P2U, #11988, Cell Signaling), Rabbit anti-NOTCH1 (1:50, Abcam, ab52627), Mouse IgM anti-HT2-280 (1:300, Terrace Biotech, TB-27AHT2-280), Rat anti-human SCGB1A1/CC10 (1:200, R&D system, MAB4218), Rabbit anti-KDR/VEGFR2 (1:100, Cell Signaling, 2479). Alexa Fluor-coupled secondary antibodies (1:500, Invitrogen) were incubated at room temperature for 60 min. After antibody staining, nuclei were stained with DAPI (1:1000, Sigma) and sections were embedded in RapiClear® (SUNJin Lab). Fluorescence images were acquired using a confocal microscope (Leica TCS SP5). All the images were further processed with Fiji software. For immunohistochemical staining of Jag1 was done by using Anti-human Jag1 antibody (1:100, Santa Cruz Biotechnology Inc., sc-390177). Slides were developed by using mouse IgG VECTASTAIN Elite ABC kit (Vector Laboratories). Slides were counterstained with haematoxylin.

### Statistical analysis

Statistical methods relevant to each figure are outlined in the figure legend. Statistical analyses were performed with Prism software package version 7.0 (GraphPad). *P* values were calculated using two-tailed unpaired Student’s t test or Two-way ANOVA for multiple group comparison (Fig. 2i, and 4b,e). Sample size for animal experiments was determined based upon pilot experiments. Mice cohort size was designed to be sufficient to enable accurate determination of statistical significance. No animals were excluded from the statistical analysis. Mice were randomly assigned to treatment or control groups, while ensuring inclusion criteria based on gender and age. Animal studies were not performed in a blinded fashion. The number of animals shown in each figure is indicated in the legends as *n* = *x* mice per group. Data shown are either representative of three or more independent experiments, or combined from three or more independent experiments as noted and analysed as mean ± s.e.m.

### Cell counting and image analysis

Sections included in cell scoring analysis for lung tissue were acquired using Leica TCS SP5 confocal microscope. At least six different sections under 10X magnification including at least 9 different alveolar regions from individual mice indicated in the figures per group were used. Cell counts were performed on ImageJ using the ‘Cell Counter’ plug-in and the performer was blinded to the specimen genotype and condition. At least two step sections (30µm apart) per individual well were used for quantification of organoids.

### RNA-sequencing analysis

RNA-seq were performed for global gene expression profiles in ACOs, and SCOs with or without DAPT treatment. Total RNAs were extracted using the RNeasy Plus Mini Kit following the manufacturer’s instructions. 100 ng of total RNA was used to generate RNA-seq library using NEBNext Ultra RNA Library prep kit (NEB, E7530L) according to the vendor’s protocol. Briefly, mRNAs were purified from total RNAs with Magnetic mRNA Isolation Kit (oligo(dT) beads) and fragmented. First and second strand cDNAs were synthesized subsequently. The double strand cDNAs were purified, and then the ends of cDNAs were repaired and ligated with sample-specific barcodes. RNA-seq libraries were sequenced using an Illumina NextSeq 500 machine. Single-ends reads from RNA-seq were aligned to the reference mouse genome (mm10) using STAR (v2.5.2b). The python package HTSeq (https://htseq.readthedocs.io/en/master/) was used to generate read counts for each gene. The read counts were analysed by the R package DESeq2 (v1.28.0)^58^ and regularized log transformed using the rlog function. Adjusted p-values (p.adj) for DEG were determined by Benjamini and Hochberg correction. The p.adj < 0.01 required to consider differentially expressed genes. Heat maps were generated using Java Treeview (v1.1.6r4).

### ATAC-sequencing analysis

The ATAC-seq assay was performed in two biological replicates for each sample of 50,000 FACS-purified cells. The quality of data from ATAC-seq was tested using FASTQC. The adapter sequences were contained in raw data. Therefore, NGmerge^59^ was used for adapter trimming. 150 bp paired-ends adapter trimmed reads were aligned against the mouse genome assembly (mm10) using bowtie2 (v2.3.4.1). We performed peak calling using MACS2 (v2.1.2) with default parameters for paired ends. Statistically significant differential open chromatin regions were determined using MAnorm (v1.1.4)^60^, which normalises read density levels and calculates p-values by MA plot methods. The heat maps and the Spearman correlation map of ATAC-seq signals were generated using deepTool2^61^. Genome browser images were generated using the Integrative Genomics Viewer (IGV) 2.7.2^62^ with bedGraph files processed from MACS2.

The ATAC-seq peaks were mapped to the region surrounding 20 Kb up- and 2 Kb down-stream of the TSS of all genes from refFlat file (mm10, UCSC). All assigned genomic features from one open region were used. To describe the distribution of genes, a promoter was defined as a region within 2 Kb from the TSS, a proximal promoter region was defined as a region between 2 Kb and 20 Kb upstream from the TSS. Mapping sites other than promoter, proximal promoter, exon, or intron were considered as intergenic target loci. To identify the cell-type specific enriched motifs in the differential open chromatin regions from ATAC-seq data, we performed motif enrichment analysis using the findMotifsGenome.pl program in the HOMER software (v4.11)^63^. The regions were adjusted to the same size with 200 bp centred on each differential peak.

### Gene ontology (GO) analysis

The gene ontology (GO) term enrichment analysis was performed using GREAT (v4.0.4)^64^ with mouse genome assembly (mm10), and whole genomes for background regions in default setting from ATAC-seq data. Gene Ontology tool^65, 66^ was used for a set of gene target of the differential open regions.

### Gene set enrichment analysis (GSEA)

GSEA was carried out by using the Gene Ontology term gene sets provided by the Mouse Genome Informatics website (http://www.informatics.jax.org)^67^. Entire detectable genes derived from RNA-seq were used for GSEA. We followed the standard GSEA user guide (http://www.broadinstitute.org/gsea/doc/GSEAUserGuideFrame.html).

### scRNA-seq library preparation and sequencing

For lineage-labeled cells from *Red2- Notch^N1ICD^* mice, YFP^+^CD45^−^CD31^−^EpCAM^+^ or RFP^+^CD45^−^CD31^−^EpCAM^+^ cells were sorted at day 28 post bleomycin injury (4 mice were pooled for each experiment). For non-lineage-labeled cells isolated from *Red2-Notch^N1ICD^* mice in parallel with experiment of lineage-labeled cells, we combined the cells of EpCAM^+^RFP^−^RFP^−^ and EpCAM^−^ population with a ratio of 1:1, respectively. The resulting cell suspension (∼110,000 cells each) were submitted as separate samples to be barcoded for the droplet-encapsulation single-cell RNA-seq experiments using the Chromium Controller (10X Genomics). Single cell cDNA synthesis, amplification, and sequencing libraries preparation were performed using the Single Cell 3’ Reagent Kit as per the 10x Genomics protocol. Libraries were multiplexed so that 2 libraries were sequenced per single lane of HiSeq 4000 using the following parameters: Read1: 27 cycles, i7: 9 cycles, i5: 0 cycles; Read2: 100 cycles to generate 150bp paired end reads.

### Data Availability

The datasets of RNA-seq, ATAC-seq, and scRNA-seq analysis generated during this study are available at GEO: GSE153677 (RNA-seq and ATACC-seq for organoids) and GSE154218 (scRNA-seq). Software used to analyse the data are either freely or commercially available.

### Alignment, quantification and quality control of scRNA-seq data

Droplet-based sequencing data were aligned and quantified using the Cell Ranger Single-Cell Software Suite (version 2.0.2, 10x Genomics Inc) using the *Mus musculus* genome (GRCm38) (official Cell Ranger reference, version 1.2.0). Cells were filtered by custom cutoff (more than 500 and less than 7000 detected genes, more than 2000 UMI count) to remove potential empty droplets and doublets. Downstream analysis included data normalisation, highly variable gene detection, log transformation, principal component analysis, neighbourhood graph generation and Louvain graph-based clustering, which was done by python package scanpy (version 1.3.6)^68^ using default parameters.

### Doublet exclusion

To exclude doublets from single-cell RNA sequencing data, we applied scrublet algorithm per sample to calculate scrublet-predicted doublet score per cell with following parameters: sim_doublet_ratio = 2; n_neighbours=; expected_doublet_rate= 0.1. Any cell with scrublet score > 0.7 was flagged as doublet. To propagate the doublet detection into potential false-negatives from scrublet analysis, we over-clustered the dataset (*sc.tl.louvain* function from scanpy package version 1.3.4; resolution = 20), and calculated the average doublet score within each cluster. Any cluster with averaged scrublet score > 0.6 was flagged as a doublet cluster. All remaining cell clusters were further examined to detect potential false-negatives from scrublet analysis according to the following criteria: (1) expression of marker genes from two distinct cell types which are unlikely according to prior knowledge, (2) higher number of UMI counts.

## Extended Data

**Extended Data Figure 1.**
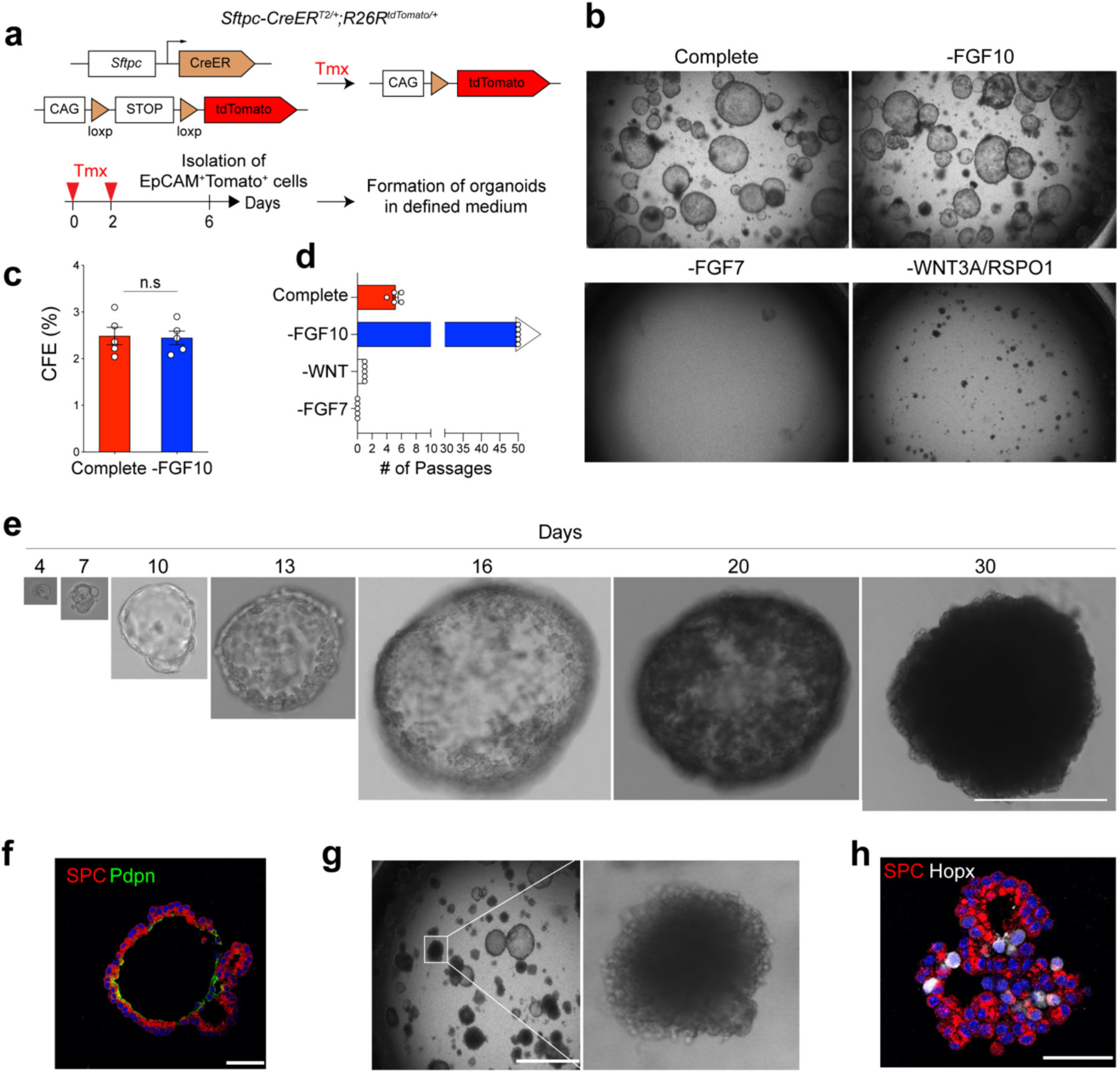
Establishment of feeder-free organoids derived from AT2 cells. **a,** Schematics of experimental design for isolation of *Sftpc* lineage-labelled AT2 cells at indicated time points after tamoxifen treatment. **b,** Representative bright-field images of organoids derived from lineage-labelled *Tomato^+^Sftpc*^+^ cells in indicated conditions; complete medium with WNT3A, RSPO1 (R-spondin 1), EGF, FGF7, FGF10, and NOG (Noggin), withdrawal of indicated factors (−FGF10, −FGF7, or −WNT3A/RSPO1). Scale bar, 2,000µm. **c, d,** Statistical quantification of colony forming efficiency (**c**) and passaging numbers (**d**) of organoids. Each individual dot represents individual biological replicate and data are presented as mean and s.e.m. n.s; not significant. **e,** Representative serial bright-field images of a lung organoid growing originated from single *Tomato^+^Sftpc*^+^ cells at the indicated time points. Magnifications: X20 (day 4 and 7), X10 (day10 and 13), and X4 (day 16, 20, and 30). Scale bars, 400µm. **f,** A representative immunofluorescence (IF) image of organoids derived from *Tomato^+^Sftpc^+^* cells at the first passage under feeder-free condition with complete culture medium. SPC (for AT2 cells, red), Pdpn (for AT1 cells, green), and DAPI (blue). Scale bars, 50µm. **g,** Representative bright-field images of organoids under feeder-free condition with complete culture medium at passage 5. Insets (left) show high-power view (right). Scale bars, 2,000µm. **h,** Representative IF images of mixed organoids cultured in complete medium at passage >5. SPC (red), Hopx (white), and DAPI (blue). Scale bars, 50µm.

**Extended Data Figure 2.**
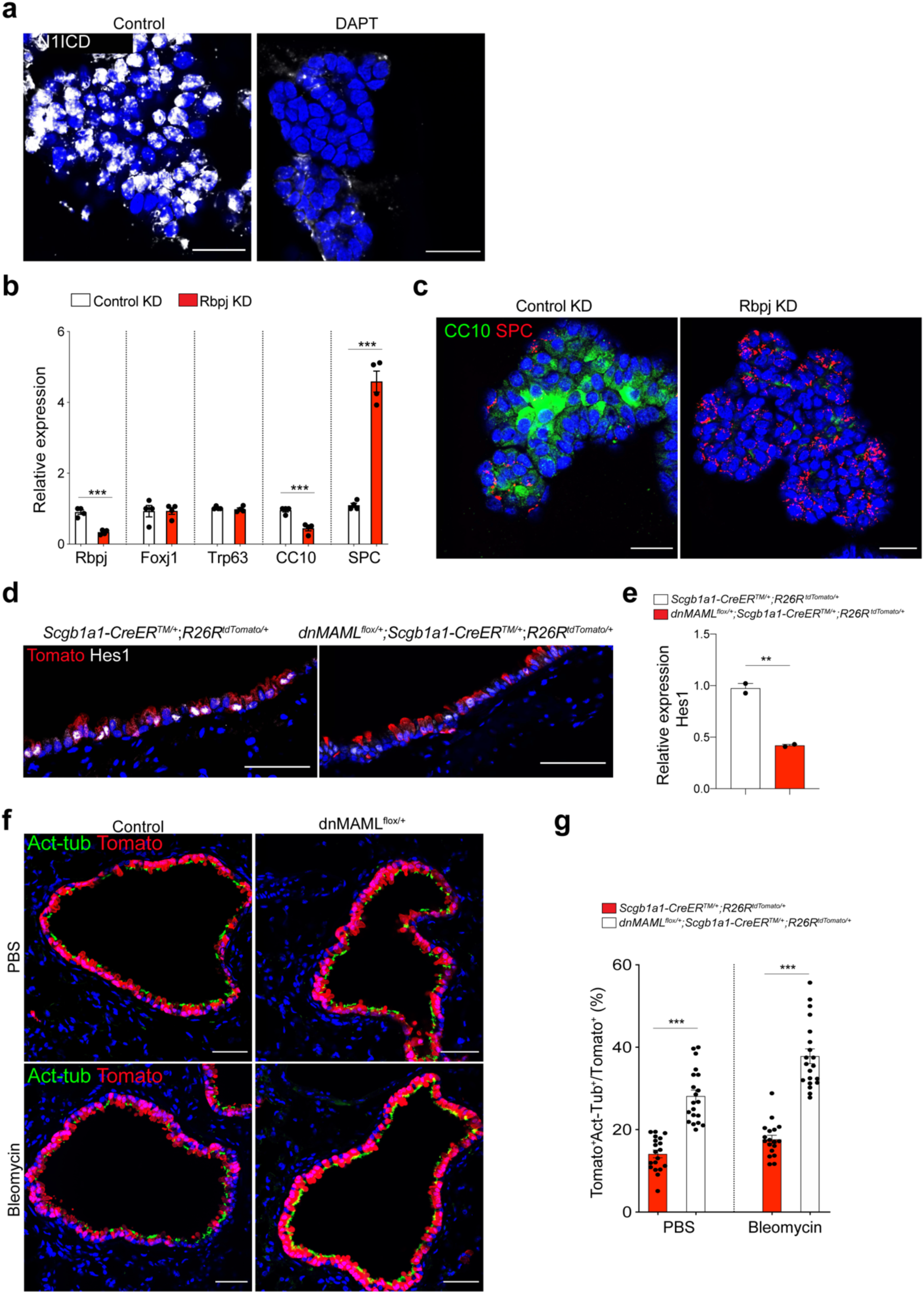
Interference of Notch activity in secretory cell-derived organoids (SCOs) enhances the differentiation into AT2 lineages. **a**, Representative IF images showing nuclear localisation of intracellular domain of Notch 1 (N1ICD) in SCOs at passage 5 with or without DAPT (20µM). CC10 (red), N1ICD (white), and DAPI (blue). Scale bars, 50µm. **b,** qPCR analysis of gene expression in control (control KD) or *Rbpj* knock-downed (*Rbpj* KD) organoids. Each individual dot represents one individual experiment (n=4 biological replicates) and data are presented as mean ± s.e.m. ***p<0.001. **c,** Representative IF images of control KD or *Rbpj* KD organoids: CC10 (green), SPC (red), and DAPI (blue). Scale bars, 50µm. **d,** Representative IF images showing Hes1 expression in lineage-labelled secretory cells after tamoxifen treatment in the indicated genotype: Tomato (for *Scgb1a1* lineage, red), Hes1 (white), and DAPI (blue). Scale bars, 50µm. **e,** qPCR analysis of *Hes1* expression in isolated EpCAM^+^Tomato^+^. Data are presented as mean ± s.e.m (n=2 individual biological replicate). **p<0.01. **f,** Representative IF images showing the derivation of lineage-labelled Act-Tub^+^ ciliated cells in PBS- or bleomycin-treated lungs of *Scgb1a1-CreER^TM/+^*;*R26R^tdTomato/+^* or *dnMAML^flox/+^*;*Scgb1a1-CreER^TM/+^*;*R26R^tdTomato/+^* mice. Tomato (for *Scgb1a1* lineage, red), Acetylated-Tubulin (Act-Tub, green), and DAPI (blue). Scale bar, 50µm. **g,** Statistical quantification of lineage-labelled ciliated cells at day 28 post PBS or bleomycin treatment. Each individual dot represents one section and data are presented as mean ± s.e.m. (n=2 biological replicates for PBS control and bleomycin). ***p<0.001.

**Extended Data Figure 3.**
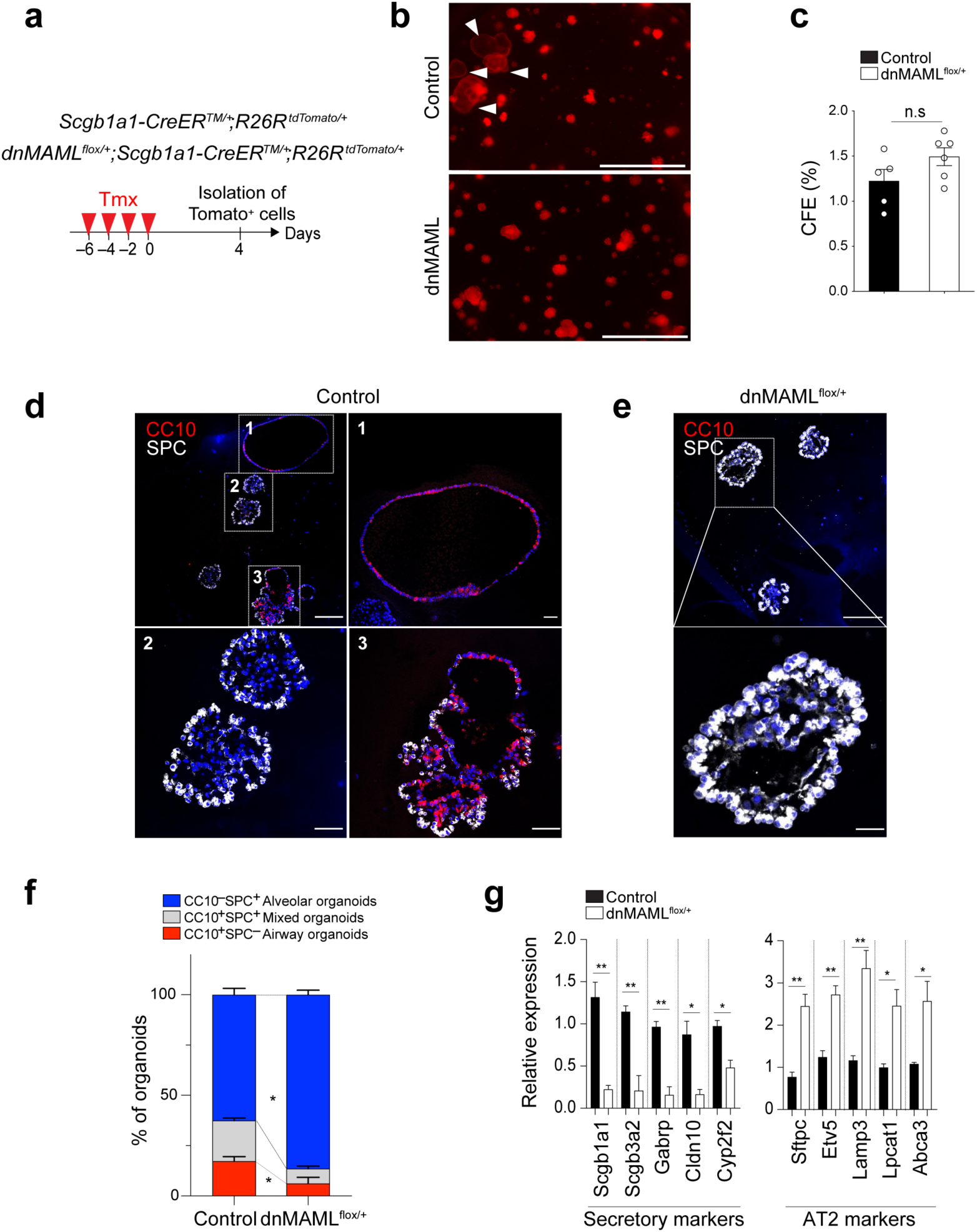
Organoid co-culture shows enhanced differentiation of secretory cells into AT2 cells by downregulation of Notch activity. **a,** Experimental design for isolation of lineage-labelled secretory cells from *Scgb1a1- CreER^TM/+^*;*R26R^tdTomato/+^* and *dnMAML^flox/+^*;*Scgb1a1-CreER^TM/+^*;*R26R^tdTomato/+^* mice at indicated time points after tamoxifen treatment. **b,** Representative fluorescent images of organoids derived from control or *dnMAML-*expressing lineage-labelled secretory cells. 5,000 cells of lineage-labelled secretory cells (Tomato^+^EpCAM^+^CD45^−^CD31^−^) isolated from *Scgb1a1-CreER^TM/+^*;*R26R^tdTomato/+^* or *dnMAML^flox/+^*;*Scgb1a1-CreER^TM/+^*;*R26R^tdTomato/+^* mouse lungs were co-cultured with stromal cells with 1:5 ratio for 14 days. Arrowheads point to cystic airway-like organoids. Scale bar, 2,000µm. **c,** Statistical quantification of colony forming efficiency of organoids. Each individual dot represents one biological replicate and data are presented as mean and s.e.m. n.s; not significant. CFE; Colony forming efficiency units. **d,** Representative IF images of three distinctive types of organoids derived from control secretory cells; Airway organoids (CC10^+^SPC^−^; denoted as 1), Alveolar organoids (CC10^−^SPC^+^; denoted as 2), and Mixed organoids (CC10^+^SPC^+^; denoted as 3). CC10 (for secretory cells, red), SPC (for AT2 cells, white), and DAPI (blue). Insets (1, 2 and 3) show high-power view. Scale bars, 50µm and 10µm (in high-power view). **e,** Representative IF images of alveolar organoids derived from *dnMAML*-expressing secretory cells. CC10 (for secretory cells, red), SPC (for AT2 cells, white), and DAPI (blue). Insets show high-power view. Scale bars, 50µm (10µm in high-power view). **f,** Quantification of each organoid types derived from *Scgb1a1- CreER^TM/+^*;*R26R^tdTomato/+^* or *dnMAML^flox/+^*;*Scgb1a1-CreER^TM/+^*;*R26R^tdTomato/+^* mice in (**d** and **e**); Airway organoids (CC10^+^SPC^−^; red), Alveolar organoids (CC10^−^SPC^+^; blue), and Mixed organoids (CC10^+^SPC^+^; grey). Data are presented as mean and s.e.m (n=3 biological replicates). *p<0.001. **g,** qPCR analysis of markers for secretory (*Scgb1a1, Scgb3a2, Gabrp, Cldn10,* and *Cyp2f2*) and AT2 (*Sftpc, Etv5, Lamp3, Lpcat1,* and *Abca3*) cells in organoids isolated from *Scgb1a1-CreER^TM/+^*;*R26R^tdTomato/+^* (black bars) and dnMAML^flox/+^;*Scgb1a1-CreER^TM/+^*; *R26R^tdTomato/+^* mice (white bars). *p<0.05, **p<0.01.

**Extended Data Figure 4.**
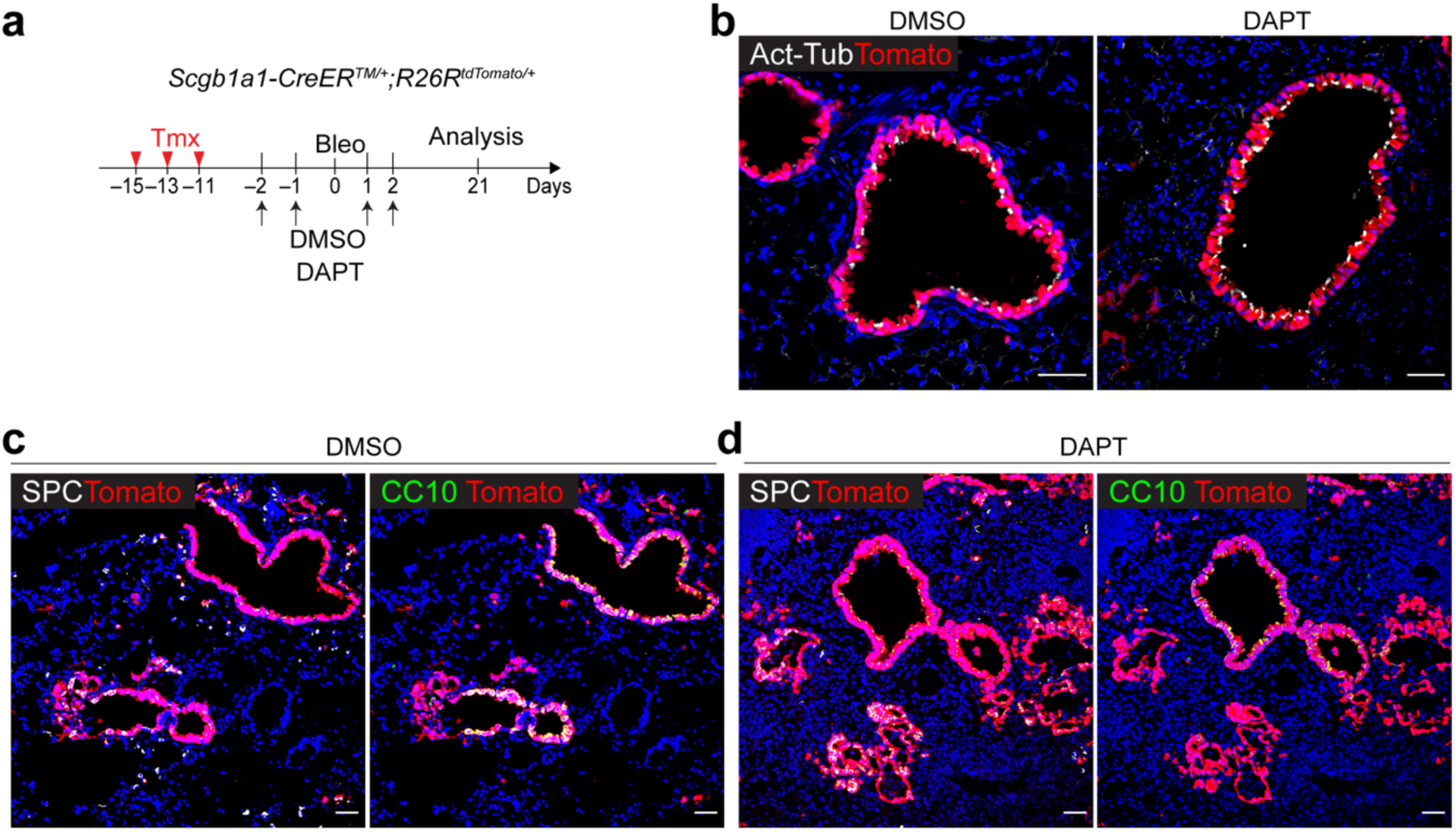
Pharmacological inhibition of Notch activity by DAPT treatment enhances the differentiation of secretory cells into AT2 cells during injury repair. **a,** Experimental design of lineage-tracing analysis of *Scgb1a1-CreER^TM/+^*;*R26R^tdTomato/+^* mice after bleomycin injury. DMSO or DAPT (50mg/kg body weight) was administrated *via* intraperitoneally as indicated time points. **b**, Representative IF images showing the increased *Scgb1a1^+^* lineage- labelled Act-Tub^+^ ciliated cells in DAPT-treated mouse lungs at day 21 post bleomycin injury compared to DMSO control mice. Tomato (for *Scgb1a1* lineage, red), Acetylated Tubulin (Act-Tub, white), and DAPI (blue). Scale bar, 50µm. **c,d**, Representative IF images showing the derivation of *Scgb1a1^+^* lineage-labelled SPC^+^ AT2 (left) or CC10^+^ secretory cells (right) in DMSO control (**c**) or DAPT-treated (**d**) mice at day 21 post bleomycin injury. Tomato (for *Scgb1a1* lineage, red), SPC (white), CC10 (green), and DAPI (blue). Scale bar, 100µm.

**Extended Data Figure 5.**
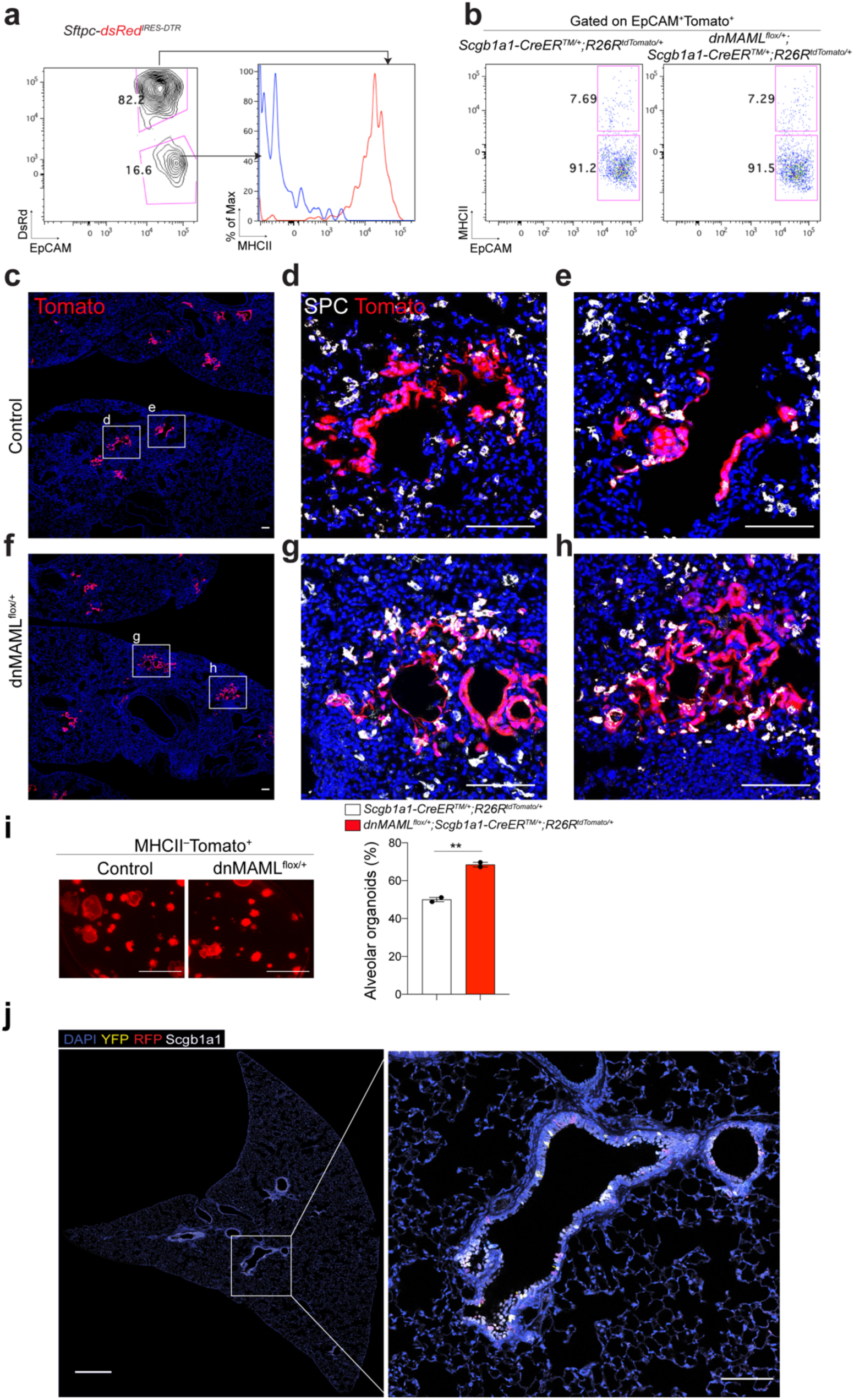
Transplantation of secretory cells, after excluding lineage- labelled AT2 cells, reveals enhanced the differentiation of secretory cells into AT2 cells via Notch inhibition. **a,** MHCII expression marks AT2 cells. Representative flow cytometry analysis of MHCII expression in SPC^+^ AT2 (EpCAM^+^dsRed^+^) or non-AT2 (EpCAM^+^dsRed^−^) cells from *Sftpc*- *dsRed^IRES-DTR^* reporter mice. Sftpc-IRES-DTR-P2A-dsRed (*Sftpc-dsRed^IRES-DTR^*) reporter mouse was used to monitor SPC-expressing AT2 cells based on the expression of dsRed (See **Methods**). Numbers adjacent to the outlined area indicate the percentage of populations. **b,** Flow cytometry analysis to exclude lineage-labelled AT2 cells, including CC10^+^SPC^+^ dual- positive cells, from *dnMAML^flox/+^*;*Scgb1a1*-*CreER*^TM/+^;*R26R*^tdTomato /+^ or *Scgb1a1*- *CreER*^TM/+^;*R26R*^tdTomato/+^ mouse lungs. **c-h,** Representative IF image of engrafted lineage- labelled secretory cells (EpCAM^+^Tomato^+^MHCII^−^) isolated from *Scgb1a1*- *CreER^TM/+^*;*R26R*^tdTomato/+^ (**c-e**) or *dnMAML^flox/+^*;*Scgb1a1*-*CreER*^TM/+^;*R26R*^tdTomato/+^ (**f-h**) mouse lungs at day 14 post transplantation. 20,000 cells of EpCAM+Tomato+MHCII^−^ were engrafted into injured lung at day 7 after bleomycin injury. Tomato (for *Scgb1a1* lineage, red), SPC (white), and DAPI (blue). Scale bars, 100µm. **i,** Representative fluorescent images of organoids (top) and statistical quantification of colony forming efficiency of alveolar organoids (left) derived from lineage-labelled AT2 cells isolated from control (*Scgb1a1*- *CreER*^TM/+^;*R26R*^tdTomato/+^) or *dnMAML^flox/+^*;*Scgb1a1-CreER^TM/+^*;*R26R^tdTomato/+^* mouse lungs. 5,000 cells of lineage-labelled secretory cells (EpCAM^+^Tomato^+^MHCII^−^) isolated from *Scgb1a1-CreER^TM/+^*;*R26R^tdTomato/+^* or *dnMAML^flox/+^*;*Scgb1a1-CreER^TM/+^*;*R26R^tdTomato/+^* mice were co-cultured with stromal cells with 1:5 ratio for 14 days. Each individual dot represents one individual experiment and data are presented as mean and s.e.m. Scale bar, 2,000µm. **j,** Representative IF images showing *Scgb1a1^+^* lineage-labelled cells in a whole lobe of *Scgb1a1- CreER^TM/+^*;*Red2-Notch^N1ICD/+^* mice post tamoxifen treatment: YFP (yellow), RFP (red), pro- SPC (white, left), Scgb1a1 (white, right) and DAPI (blue). Scale bar, 500µm (left), and 100µm (right). Of note, only airway secretory cells were labelled by tamoxifen induction.

**Extended Data Figure 6.**
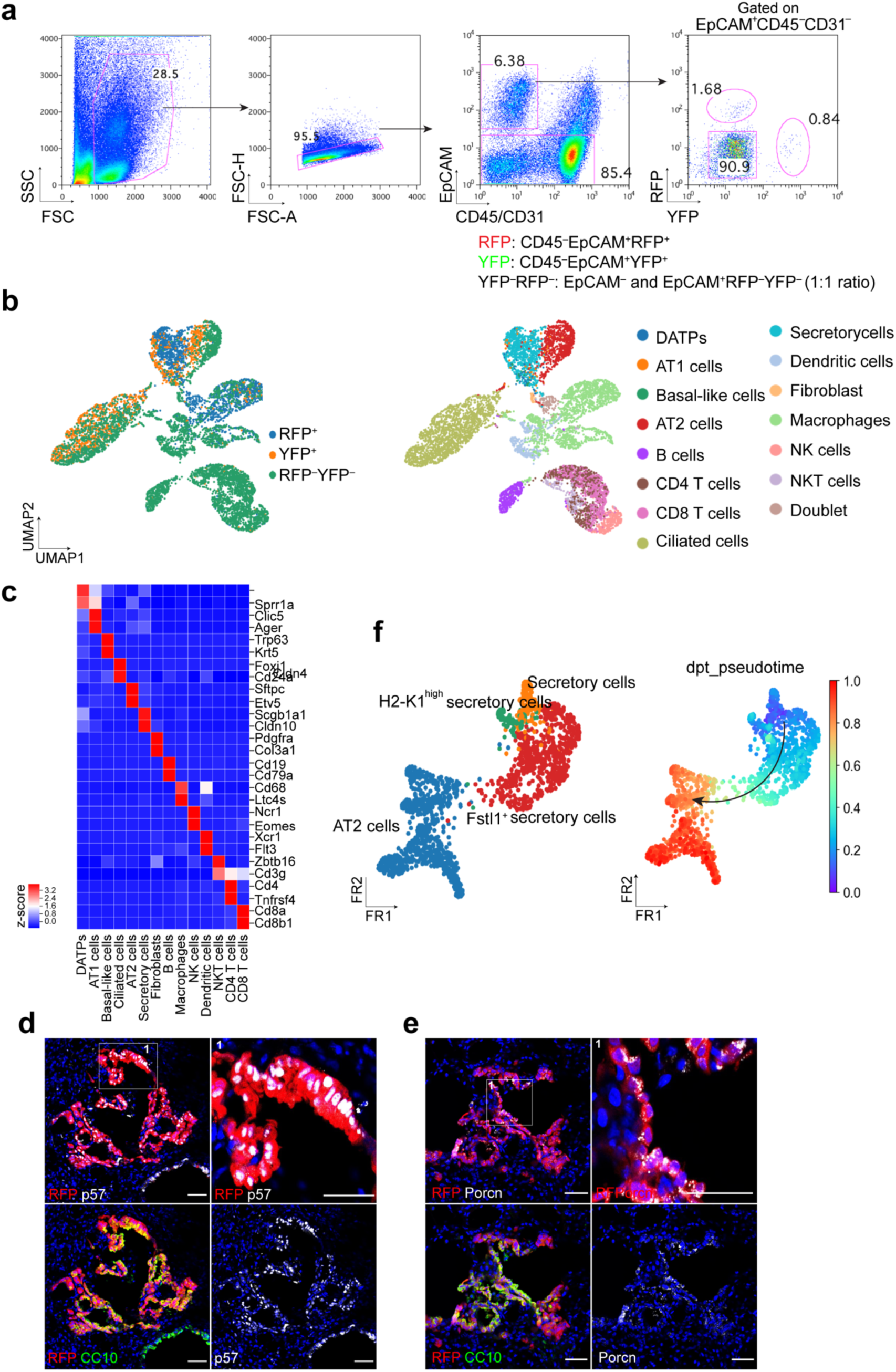
scRNA-seq analysis of lung cells from *Scgb1a1-CreER^TM/+^*;*Red2- Notch^N1ICD/+^* mice at day 28 post bleomycin injury. **a,** Sorting strategy for *Scgb1a1* lineage-labelled RFP^+^ and YFP^+^ or unlabelled (YFP^−^RFP^−^) cells by flow cytometry after bleomycin injury. For unlabelled population (YFP^−^RFP^−^), EpCAM^−^RFP^−^YFP^−^ (non-epithelial population) and EpCAM^+^RFP^−^YFP^−^ (epithelial population) cells were mixed at 1:1 ratio. **b,** Clusters of cell population after bleomycin injury from 10x Genomics 3’ scRNA-seq analysis visualised by UMAP, assigned by specific colours. **c,** A heatmap showing gene expression patterns of key markers in each distinctive cell cluster. **d, e,** Representative IF images showing the expansion of Porcn^+^ or p57^+^ cells derivation from RFP^+^ cells at day 28 post bleomycin injury. RFP (red), CC10 (green), p57 (white, e), Porcn (white, f), and DAPI (blue). Insets (1) show high-power view. Scale bar, 50µm. **f,** Diffusion map according to diffusion pseudotime (DPT, right) order coloured by samples (left).

**Extended Data Figure 7.**
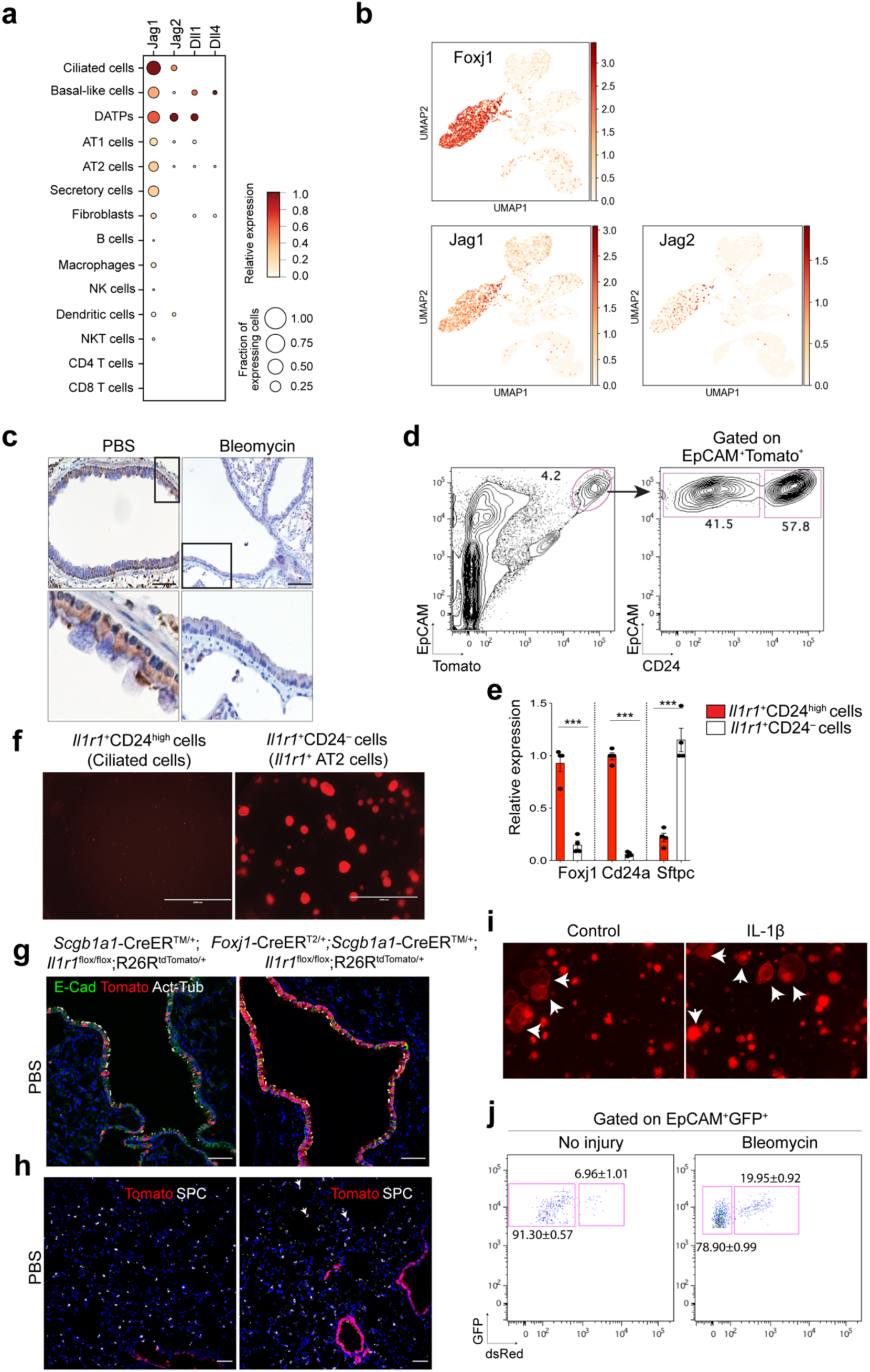
Expression of Notch ligands in ciliated cells post alveolar injury. **a,** Expression of Notch ligand in each distinctive cluster revealed in **Extended Data Fig. 6**. The size of the circle represents the fraction of cells expressing the gene and the colour represents the relative expression of each gene. **b,** UMAP visualisation of the log-transformed (log10(TPM+1)), normalised expression of selected marker genes (*Jag1* and *Jag2* for Notch ligand expression; *Foxj1* for ciliated cells) in distinctive clusters. **c,** Representative IHC images stained with anti-Jag1 in the distal airway of lungs from *Scgb1a1-CreER^TM/+^*;*R26R^tdTomato^*^/+^ mice at day 14 post PBS or bleomycin treatment. Insets (top) show high-power view (bottom). Scale bars, 50µm. **d,** Sorting strategy for *Il1r1^+^* lineage-labelled ciliated (EpCAM^+^Tomato^+^CD24^high^) and AT2 (EpCAM^+^Tomato^+^CD24^−^) cells from *Il1r1-CreER^T2/+^*; *R26R^tdTomato/+^* mouse lungs at day 4 post tamoxifen injection. **e,** qPCR analysis of *Foxj1*, *Cd24, and Sftpc* expression in isolated cells in (**d**). Each individual dot represents one individual experiment and data are presented as mean ± s.e.m. ***p<0.001. **f,** Representative fluorescent images of organoids derived from *Il1r1* lineage-labelled ciliated (*Il1r1*^+^CD24^high^) or AT2 (*Il1r1*^+^CD24^−^) cells. Ciliated cells are unable to form organoids. Scale bars, 2,000µm. **g,** Representative IF images showing intact ciliated cells in the lungs of *Il1r1^flox/flox^*;*Foxj1- CreER^T2/+^*;*Scgb1a1-CreER^TM/+^*;*R26R^tdTomato/+^* mice compared to those in the lungs of control *Il1r1^flox/flox^*;*Scgb1a1-CreER^TM/+^*;*R26^RtdTomato/+^* mice: Tomato (for *Scgb1a1* or *Foxj1* lineage, red), Acetylated Tubulin (Act-Tub, white), E-Cadherin (E-Cad, green), and DAPI (blue). Scale bars, 50µm. **h**, Representative IF images showing a small number of *Scgb1a1* lineage-labelled AT2 cells at day 28 post PBS treatment in *Scgb1a1-CreER^TM/+^*;*Il1r1^flox/flox^*;*R26R^tdTomato/+^* or *Foxj1-CreER^T2/+^*;*Scgb1a1-CreER^TM/+^*;*Il1r1^flox/flox^*;*R26R^tdTomato/+^* mice. Tomato (for *Scgb1a1* lineage, red), SPC (white), and DAPI (blue). Arrowheads point to lineage-labelled SPC^+^ AT2 cells. Scale bar, 100µm. **i,** Representative fluorescent images of organoids derived from lineage-labelled secretory cells of *Scgb1a1-CreER^TM/+^*;*R26R^tdTomato/+^* lungs. Organoids were cultured with PBS or IL-1β for 14 days. Arrows point to cystic airway-like organoids. Scale bar, 2,000µm. **j,** Flow cytometry analysis of secretory cell-derived AT2 cells isolated from *Scgb1a1-CreER^TM/+^*;*R26R^fGFP/+^*;*Sftpc-dsRed^IRES-DTR/+^* mouse lungs after PBS (left) or bleomycin (right) treatment. Data are presented as mean ± s.e.m (n=2 individual biological replicate).

**Extended Data Figure 8.**
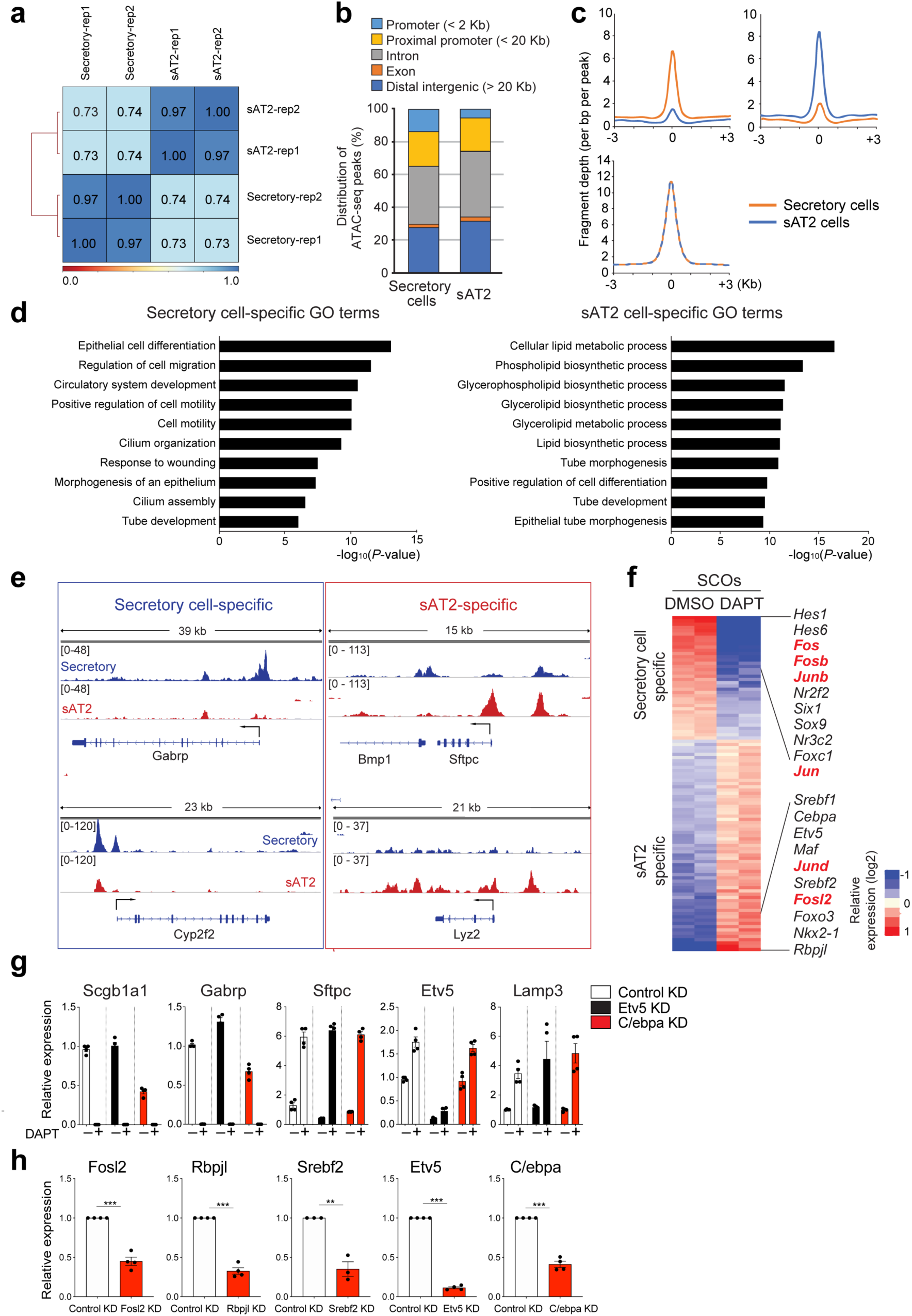
Differential genetic and epigenetic characteristics between secretory cells and secretory-derived AT2 (sAT2) cells. **a,** A spearman correlation map plotted by using ATAC-seq replicates obtained from secretory and sAT2 cells. **b**, A bar graph showing the chromosomal distribution of secretory-specific and sAT2-specific cis-regulatory elements, mapped by ATAC-seq in secretory and sAT2 cells across the mouse genome. **c,** The distribution of ATAC-seq peak signals in secretory and sAT2 cells near the centre of secretory-specific, shared, and sAT2-specific open regions. **d**, Bar graphs showing enriched GO terms of the genes nearby secretory and sAT2-specific ATAC- seq peaks, respectively. **e,** Signal track images showing open chromatin regions nearby markers for secretory (*Gabrp* and *Cyp2f2*) and AT2 cells (*Sftpc* and *Lyz2*) in secretory (blue) and sAT2 cells (red). **f,** A heatmap showing differentially expressed transcription factors (TFs) in secretory-derived organoids with or without DAPT treatment. **g,** qPCR analysis of the markers for secretory (*Scgb1a1* and *Gabrp*) and AT2 cells (*Sftpc*, *Etv5*, and *Lamp3*) in secretory cell- derived organoids (SCOs) with or without DAPT treatment after knockdown of control (white), *Etv5* (black), or *Cebpa* (red). **h,** qPCR analysis to confirm the knockdown efficiency of the genes that are used for indicated constructs in SCOs. Each individual dot represents one individual biological experiment (n=4 for knockdown of Fosl2, Rpbjl, Etv5, and C/ebpa, n=3 for knockdown of Srebf2) and data are presented as mean ± s.e.m. **p<0.01, ***p<0.001.

**Extended Data Figure 9.**
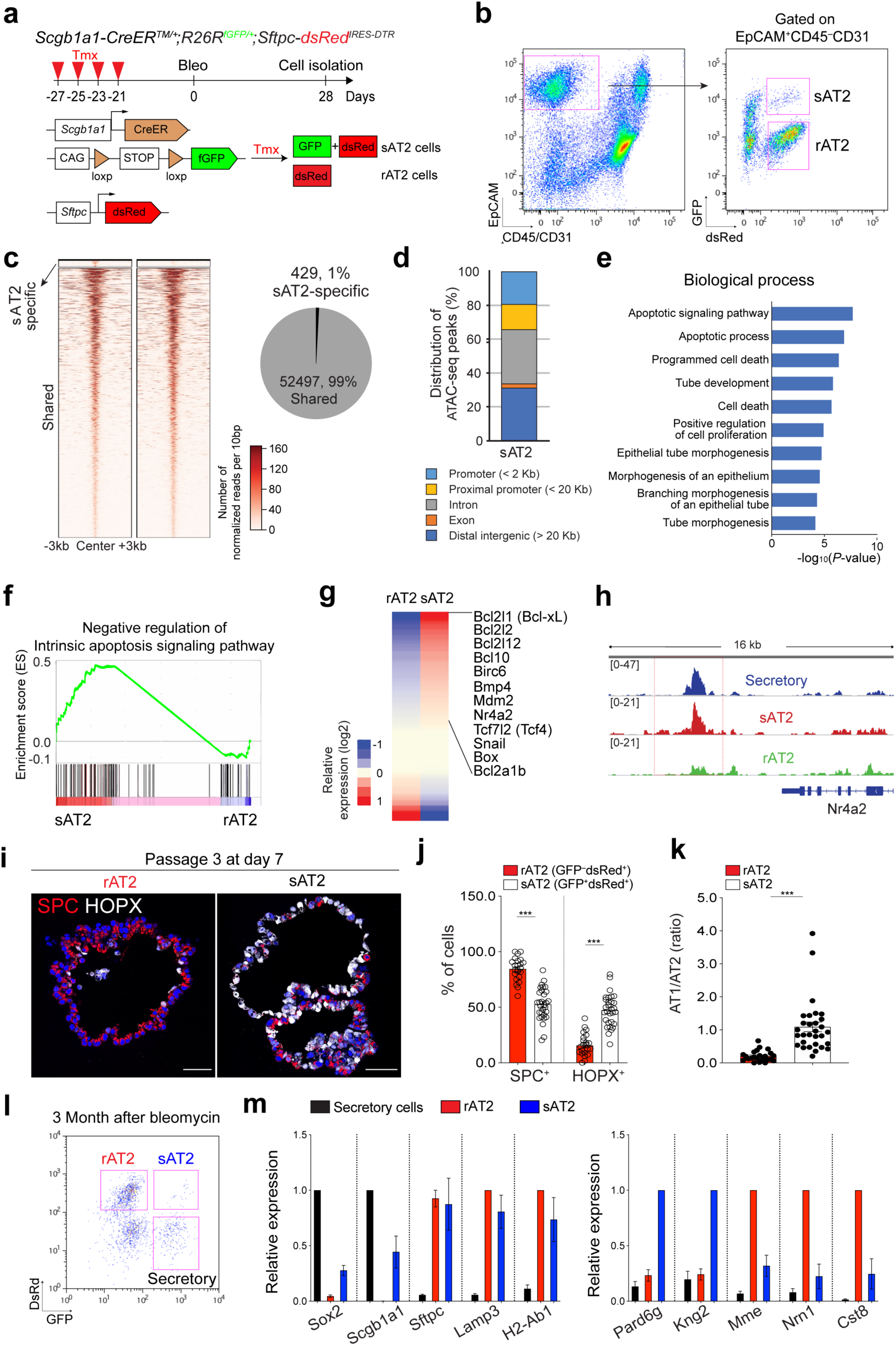
Genetic and epigenetic differences of sAT2 cells compared to rAT2 cells. **a,** Experimental design for isolation of secretory-derived (sAT2, GFP^+^dsRed^+^) and resident AT2 cells (non-lineage-labelled AT2, rAT2, GFP^−^dsRed^+^) using *Scgb1a1-CreER^TM/+^*; *R26R^fGFP/+^*;*Sftpc-dsRed^IRES-DTR/+^* mice. Specific time points for tamoxifen injection and isolation are indicated. **b,** Representative flow cytometry analysis for isolation of sAT2 (GFP^+^dsRed^+^) and rAT2 (GFP^−^dsRed^+^) cells. **c,** Heatmaps showing open chromatin regions specific in sAT2 cells and shared open chromatin regions between sAT2 cells and rAT2 cells (left). A pie chart presenting the proportion of sAT2-specific (blue) and shared regions (grey) (right). **d,** A bar graph showing the chromosomal distribution of sAT2-specific ATAC-seq peaks mapped in (**c**). **e,** A bar graph showing enriched GO terms of the genes nearby sAT2- specific peaks. **f,** GSEA with gene sets representing the negative regulation of intrinsic apoptosis signalling pathway in scRNA-seq data of sAT2 and rAT2 shown in Fig. 6a. **g,** A heatmap showing the expression patterns of the genes belonging to the negative regulation of apoptosis signalling pathway in sAT2 and rAT2 cells monitored by scRNA-seq (Fig. 6a). **h,** Signal track image showing open regions for *Nr4a2*, anti-apoptotic pathway marker, in secretory (blue), sAT2 (red), and rAT2 cells (green). **i,** Representative IF images of organoids derived from rAT2 or sAT2 cells. SPC (for AT2 cells, red), HOPX (for AT1 cells, white), and DAPI (blue). Scale bars, 50µm. **j, k,** Quantification of the frequency of AT2 (SPC^+^) or AT1 (HOPX^+^) cells (**b**) and the ratio of AT1/AT2 cells (**c)** in rAT2- or sAT2-derived organoids. Each individual dot represents one organoid and data are presented as mean ± s.e.m (n=2 independent biological replicates). ***p<0.001. **l**, Flow cytometry analysis of *Scgb1a1^+^* lineage-labelled AT2 cells isolated from *Scgb1a1-CreER^T2/+^*;*R26R^fGFP/+^*;*Sftpc-dsRed^IRES-DTR/+^* mouse lungs at 3 months after bleomycin injury. **m**, qPCR analysis of the genes in isolated secretory, rAT2, or sAT2 cells. Data are presented as mean ± s.e.m (n=4 individual mice).

**Extended Data Figure 10.**
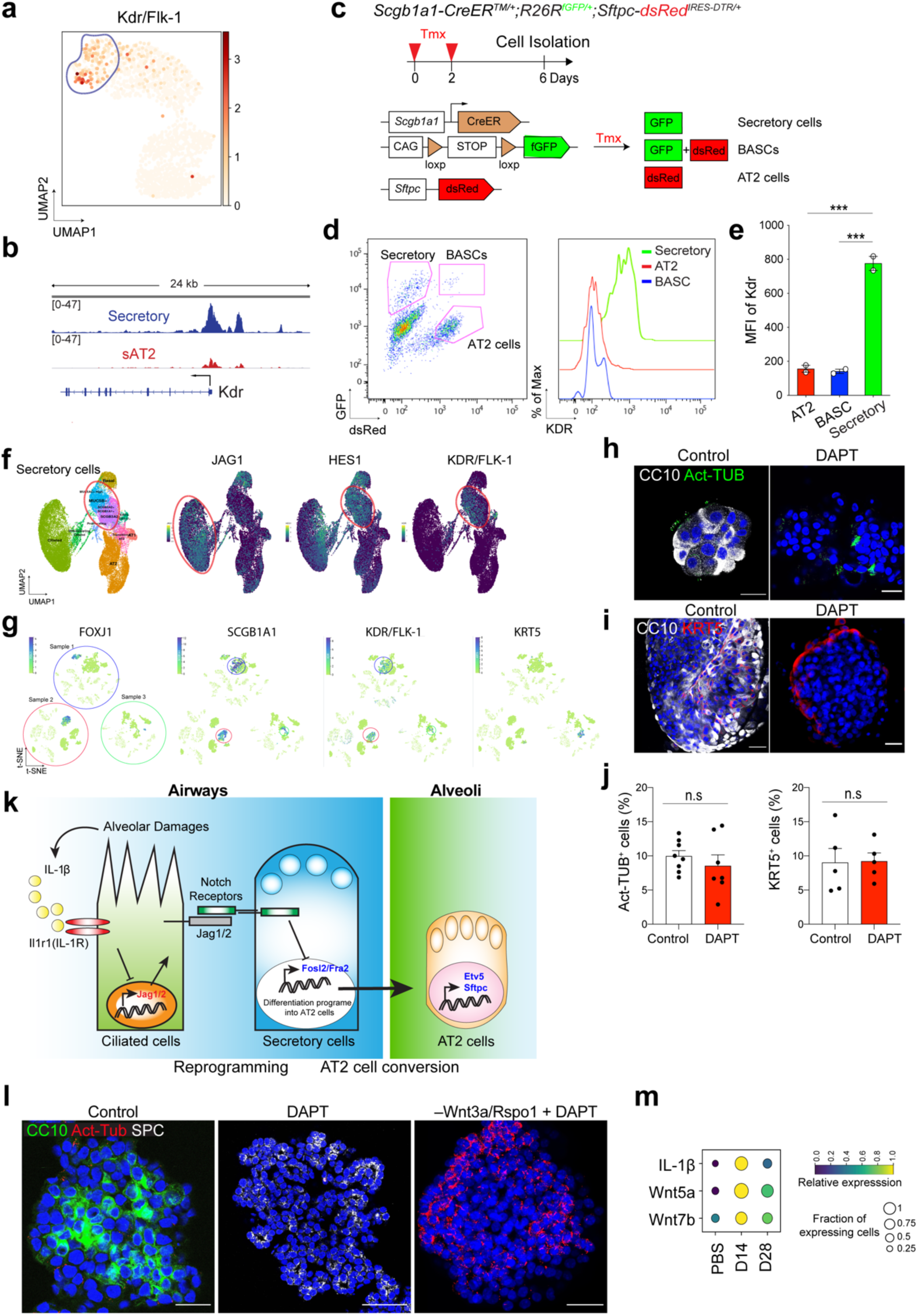
Identification of KDR/Flk-1 as a new surface marker of secretory cells in both mouse and human lungs. **a,** UMAP visualisation of the log-transformed (log10(TPM+1)), normalised expression of Kdr/Flk-1 in distinctive clusters shown in Fig. 3j. **b,** A signal track image showing open regions nearby *Kdr* mapped in secretory (blue) and sAT2 cells (red). **c,** Experimental design for isolation of CC10^+^SPC^−^ secretory cells, CC10^+^SPC^+^ dual-positive cells including Bronchioalveolar stem cells (BASCs), and CC10^−^SPC^+^ AT2 cells from *Scgb1a1- CreER^TM/+^*;*R26R^fGFP/+^*;*Sftpc-dsRed^IRES-DTR/+^* mice. Specific time points for tamoxifen injection and isolation are indicated. **d,** Representative flow cytometry analysis (left) for isolation of secretory (GFP^+^dsRed^−^), BASCs (GFP^+^dsRed^+^), and AT2 (GFP^−^dsRed^+^) cells. KDR expression was determined by flow cytometry (right). **e,** A bar graph showing MFI (mean fluorescence of intensity) of KDR expression in indicated populations. Each individual dot represents one individual experiment and data are presented as mean ± s.e.m. ***p<0.001. **f,g,** UMAP (**f**) or t-SNE (**g**) visualisation of the normalised expression of *JAG1, HES1*, and *KDR* in distinctive human lung cell clusters from the dataset of Habermann et al.,^44^(**f**) or Travaglini et al.,^45^(**g**). *JAG1* is highly expressed in ciliated cells. Secretory cells express higher expression levels of *HES1* and *KDR*. **h**,**i**, Representative IF images of secretory cell-derived organoids retaining Act-Tub+ ciliated cells (**h**) and KRT5+ basal cells (**i**) treated with DMSO or DAPT (20µM). CC10 (white), Acetylated Tubulin (Act-Tub, green, **h**), KRT5 (red, **i**), and DAPI (blue). Scale bar, 50µm. **j**, Quantification of the frequency of Act-Tub^+^ ciliated cells or KRT5^+^ basal cells in control or DAPT-treated organoids. Each individual dot represents one organoid and data are presented as mean ± s.e.m (n=3 independent experiments). n.s.=not significance. **k**, Notch inhibition coordinates the cell fate decision of secretory cells into AT2 cells in two stages: 1) Reprogramming; loss of secretory cell identity and 2) AT2 cell conversion; acquisition of transcriptional programmes for AT2 cell differentiation vis the transcription regulator *Fosl2*. Pro-inflammatory cytokine IL-1β driven by tissue injury regulates the expression of Notch ligands such as Jag1/2 in ciliated cells, which induces the downregulation of Notch activity in secretory cells localised adjacent to ciliated cells. Fosl2/Fra2-mediated AP- 1 factor triggers the differentiation programmes of secretory cells to enter AT2 cell differentiation process during alveolar regeneration. **l,** Representative IF images of secretory organoids treated with DMSO (control) or DAPT (20µM) in the presence (middle) or withdrawals of Wnt-inducing factors (right). Organoids were cultured in the medium supplemented with Wnt3a/Rspo1 for 7 days and then DAPT was treated for further 7 days with or without Wnt3a/Rspo1. CC10 (for secretory cells, green), Acetylated Tubulin (for ciliated cells, Act-Tub, red), SPC (for AT2 cells, white), and DAPI (blue). Scale bars, 50µm. **m,** The expression of *IL-1β*, *Wnt5a*, and *Wnt7b* at indicated time points after bleomycin injury. Data were from our previous study^30^.

## References

1. Fuchs, E., Tumbar, T. & Guasch, G. Socializing with the neighbors: stem cells and their niche. Cell 116, 769–778, doi:10.1016/s0092-8674(04)00255-7 (2004).

2. Blanpain, C. & Fuchs, E. Stem cell plasticity. Plasticity of epithelial stem cells in tissue regeneration. Science 344, 1242281, doi:10.1126/science.1242281 (2014).

3. Butler, J. P. et al. Evidence for adult lung growth in humans. N Engl J Med 367, 244–247, doi:10.1056/NEJMoa1203983 (2012).

4. Hogan, B. L. et al. Repair and regeneration of the respiratory system: complexity, plasticity, and mechanisms of lung stem cell function. Cell Stem Cell 15, 123–138, doi:10.1016/j.stem.2014.07.012 (2014).

5. Barkauskas, C. E. et al. Type 2 alveolar cells are stem cells in adult lung. J Clin Invest 123, 3025–3036, doi:10.1172/JCI68782 (2013).

6. Rock, J. R. et al. Multiple stromal populations contribute to pulmonary fibrosis without evidence for epithelial to mesenchymal transition. Proc Natl Acad Sci U S A 108, E1475–1483, doi:10.1073/pnas.1117988108 (2011).

7. Adamson, I. Y. & Bowden, D. H. The type 2 cell as progenitor of alveolar epithelial regeneration. A cytodynamic study in mice after exposure to oxygen. Lab Invest 30, 35–42 (1974).

8. Kathiriya, J. J., Brumwell, A. N., Jackson, J. R., Tang, X. & Chapman, H. A. Distinct Airway Epithelial Stem Cells Hide among Club Cells but Mobilize to Promote Alveolar Regeneration. Cell Stem Cell 26, 346–358 e344, doi:10.1016/j.stem.2019.12.014 (2020).

9. Vaughan, A. E. et al. Lineage-negative progenitors mobilize to regenerate lung epithelium after major injury. Nature 517, 621–625, doi:10.1038/nature14112 (2015).

10. Salwig, I. et al. Bronchioalveolar stem cells are a main source for regeneration of distal lung epithelia in vivo. EMBO J 38, doi:10.15252/embj.2019102099 (2019).

11. Liu, Q. et al. Lung regeneration by multipotent stem cells residing at the bronchioalveolar-duct junction. Nat Genet 51, 728–738, doi:10.1038/s41588-019-0346-6 (2019).

12. Guha, A., Deshpande, A., Jain, A., Sebastiani, P. & Cardoso, W. V. Uroplakin 3a(+) Cells Are a Distinctive Population of Epithelial Progenitors that Contribute to Airway Maintenance and Post-injury Repair. Cell Rep 19, 246–254, doi:10.1016/j.celrep.2017.03.051 (2017).

13. Rawlins, E. L. et al. The role of Scgb1a1+ Clara cells in the long-term maintenance and repair of lung airway, but not alveolar, epithelium. Cell Stem Cell 4, 525–534, doi:10.1016/j.stem.2009.04.002 (2009).

14. Miller, A. J. et al. Generation of lung organoids from human pluripotent stem cells in vitro. Nat Protoc 14, 518–540, doi:10.1038/s41596-018-0104-8 (2019).

15. Huch, M. et al. Unlimited in vitro expansion of adult bi-potent pancreas progenitors through the Lgr5/R-spondin axis. EMBO J 32, 2708–2721, doi:10.1038/emboj.2013.204 (2013).

16. Huch, M. et al. In vitro expansion of single Lgr5+ liver stem cells induced by Wnt- driven regeneration. Nature 494, 247–250, doi:10.1038/nature11826 (2013).

17. Nikolic, M. Z. et al. Human embryonic lung epithelial tips are multipotent progenitors that can be expanded in vitro as long-term self-renewing organoids. Elife 6, doi:10.7554/eLife.26575 (2017).

18. Sachs, N. et al. Long-term expanding human airway organoids for disease modeling. EMBO J 38, doi:10.15252/embj.2018100300 (2019).

19. Lee, J. H. et al. Anatomically and Functionally Distinct Lung Mesenchymal Populations Marked by Lgr5 and Lgr6. Cell 170, 1149–1163 e1112, doi:10.1016/j.cell.2017.07.028 (2017).

20. Shiraishi, K. et al. In vitro expansion of endogenous human alveolar epithelial type II cells in fibroblast-free spheroid culture. Biochem Biophys Res Commun 515, 579–585, doi:10.1016/j.bbrc.2019.05.187 (2019).

21. Weiner, A. I. et al. Mesenchyme-free expansion and transplantation of adult alveolar progenitor cells: steps toward cell-based regenerative therapies. NPJ Regen Med 4, 17, doi:10.1038/s41536-019-0080-9 (2019).

22. Katsura, H. et al. Human Lung Stem Cell-Based Alveolospheres Provide Insights into SARS-CoV-2-Mediated Interferon Responses and Pneumocyte Dysfunction. Cell Stem Cell 27, 890–904 e898, doi:10.1016/j.stem.2020.10.005 (2020).

23. Youk, J. et al. Three-Dimensional Human Alveolar Stem Cell Culture Models Reveal Infection Response to SARS-CoV-2. Cell Stem Cell 27, 905–919 e910, doi:10.1016/j.stem.2020.10.004 (2020).

24. Maillard, I. et al. The requirement for Notch signaling at the beta-selection checkpoint in vivo is absolute and independent of the pre-T cell receptor. J Exp Med 203, 2239–2245, doi:10.1084/jem.20061020 (2006).

25. Pardo-Saganta, A. et al. Parent stem cells can serve as niches for their daughter cells. Nature 523, 597–601, doi:10.1038/nature14553 (2015).

26. Lafkas, D. et al. Therapeutic antibodies reveal Notch control of transdifferentiation in the adult lung. Nature 528, 127–131, doi:10.1038/nature15715 (2015).

27. Morimoto, M. et al. Canonical Notch signaling in the developing lung is required for determination of arterial smooth muscle cells and selection of Clara versus ciliated cell fate. J Cell Sci 123, 213–224, doi:10.1242/jcs.058669 (2010).

28. You, P. et al. Jagged-1-HES-1 signaling inhibits the differentiation of TH17 cells via ROR gammat. J Biol Regul Homeost Agents 27, 79–93 (2013).

29. Tsao, P. N. et al. Notch signaling controls the balance of ciliated and secretory cell fates in developing airways. Development 136, 2297–2307, doi:10.1242/dev.034884 (2009).

30. Choi, J. et al. Inflammatory Signals Induce AT2 Cell-Derived Damage-Associated Transient Progenitors that Mediate Alveolar Regeneration. Cell Stem Cell 27, 366–382 e367, doi:10.1016/j.stem.2020.06.020 (2020).

31. Rawlins, E. L., Ostrowski, L. E., Randell, S. H. & Hogan, B. L. Lung development and repair: contribution of the ciliated lineage. Proc Natl Acad Sci U S A 104, 410–417, doi:10.1073/pnas.0610770104 (2007).

32. Kotton, D. N. & Morrisey, E. E. Lung regeneration: mechanisms, applications and emerging stem cell populations. Nat Med 20, 822–832, doi:10.1038/nm.3642 (2014).

33. Kimura, S. et al. The T/ebp null mouse: thyroid-specific enhancer-binding protein is essential for the organogenesis of the thyroid, lung, ventral forebrain, and pituitary. Genes Dev 10, 60–69, doi:10.1101/gad.10.1.60 (1996).

34. Martis, P. C. et al. C/EBPalpha is required for lung maturation at birth. Development 133, 1155–1164, doi:10.1242/dev.02273 (2006).

35. Zhang, Z. et al. Transcription factor Etv5 is essential for the maintenance of alveolar type II cells. Proc Natl Acad Sci U S A 114, 3903–3908, doi:10.1073/pnas.1621177114 (2017).

36. Didon, L., Roos, A. B., Elmberger, G. P., Gonzalez, F. J. & Nord, M. Lung-specific inactivation of CCAAT/enhancer binding protein alpha causes a pathological pattern characteristic of COPD. Eur Respir J 35, 186–197, doi:10.1183/09031936.00185008 (2010).

37. Holla, V. R., Mann, J. R., Shi, Q. & DuBois, R. N. Prostaglandin E2 regulates the nuclear receptor NR4A2 in colorectal cancer. J Biol Chem 281, 2676–2682, doi:10.1074/jbc.M507752200 (2006).

38. Ke, N. et al. Nuclear hormone receptor NR4A2 is involved in cell transformation and apoptosis. Cancer Res 64, 8208–8212, doi:10.1158/0008-5472.CAN-04-2134 (2004).

39. Koppula, P., Zhang, Y., Zhuang, L. & Gan, B. Amino acid transporter SLC7A11/xCT at the crossroads of regulating redox homeostasis and nutrient dependency of cancer. Cancer Commun (Lond) 38, 12, doi:10.1186/s40880-018-0288-x (2018).

40. Dixon, S. J. et al. Ferroptosis: an iron-dependent form of nonapoptotic cell death. Cell 149, 1060–1072, doi:10.1016/j.cell.2012.03.042 (2012).

41. Lee, J. H. et al. Lung stem cell differentiation in mice directed by endothelial cells via a BMP4-NFATc1-thrombospondin-1 axis. Cell 156, 440–455, doi:10.1016/j.cell.2013.12.039 (2014).

42. Chen, H. et al. Airway epithelial progenitors are region specific and show differential responses to bleomycin-induced lung injury. Stem Cells 30, 1948–1960, doi:10.1002/stem.1150 (2012).

43. Chernaya, O., Shinin, V., Liu, Y. & Minshall, R. D. Behavioral heterogeneity of adult mouse lung epithelial progenitor cells. Stem Cells Dev 23, 2744–2757, doi:10.1089/scd.2013.0631 (2014).

44. Habermann, A. C. et al. Single-cell RNA sequencing reveals profibrotic roles of distinct epithelial and mesenchymal lineages in pulmonary fibrosis. Sci Adv 6, eaba1972, doi:10.1126/sciadv.aba1972 (2020).

45. Travaglini, K. J. et al. A molecular cell atlas of the human lung from single-cell RNA sequencing. Nature 587, 619–625, doi:10.1038/s41586-020-2922-4 (2020).

46. Geng, Y. et al. Follistatin-like 1 (Fstl1) is a bone morphogenetic protein (BMP) 4 signaling antagonist in controlling mouse lung development. Proc Natl Acad Sci U S A 108, 7058–7063, doi:10.1073/pnas.1007293108 (2011).

47. Weaver, M., Yingling, J. M., Dunn, N. R., Bellusci, S. & Hogan, B. L. Bmp signaling regulates proximal-distal differentiation of endoderm in mouse lung development. Development 126, 4005–4015 (1999).

48. Eming, S. A., Wynn, T. A. & Martin, P. Inflammation and metabolism in tissue repair and regeneration. Science 356, 1026–1030, doi:10.1126/science.aam7928 (2017).

49. Naik, S. et al. Inflammatory memory sensitizes skin epithelial stem cells to tissue damage. Nature 550, 475–480, doi:10.1038/nature24271 (2017).

50. Karin, M. & Clevers, H. Reparative inflammation takes charge of tissue regeneration. Nature 529, 307–315, doi:10.1038/nature17039 (2016).

51. Rouillard, A. D. et al. The harmonizome: a collection of processed datasets gathered to serve and mine knowledge about genes and proteins. Database (Oxford) 2016, doi:10.1093/database/baw100 (2016).

52. Zepp, J. A. et al. Distinct Mesenchymal Lineages and Niches Promote Epithelial Self- Renewal and Myofibrogenesis in the Lung. Cell 170, 1134–1148 e1110, doi:10.1016/j.cell.2017.07.034 (2017).

53. Chilosi, M. et al. Aberrant Wnt/beta-catenin pathway activation in idiopathic pulmonary fibrosis. Am J Pathol 162, 1495–1502, doi:10.1016/s0002-9440(10)64282-4 (2003).

54. Jensen-Taubman, S. M., Steinberg, S. M. & Linnoila, R. I. Bronchiolization of the alveoli in lung cancer: pathology, patterns of differentiation and oncogene expression. Int J Cancer 75, 489–496, doi:10.1002/(sici)1097-0215(19980209)75:4<489::aid-ijc1>3.0.co;2-p (1998).

55. Xu, Y. et al. Single-cell RNA sequencing identifies diverse roles of epithelial cells in idiopathic pulmonary fibrosis. JCI Insight 1, e90558, doi:10.1172/jci.insight.90558 (2016).

56. Madisen, L. et al. A robust and high-throughput Cre reporting and characterization system for the whole mouse brain. Nat Neurosci 13, 133–140, doi:10.1038/nn.2467 (2010).

57. Robson, M. J. et al. Generation and Characterization of Mice Expressing a Conditional Allele of the Interleukin-1 Receptor Type 1. PLoS One 11, e0150068, doi:10.1371/journal.pone.0150068 (2016).

58. Love, M. I., Huber, W. & Anders, S. Moderated estimation of fold change and dispersion for RNA-seq data with DESeq2. Genome Biol 15, 550, doi:10.1186/s13059-014-0550-8 (2014).

59. Gaspar, J. M. NGmerge: merging paired-end reads via novel empirically-derived models of sequencing errors. BMC bioinformatics 19, 1–9 (2018).

60. Shao, Z., Zhang, Y., Yuan, G. C., Orkin, S. H. & Waxman, D. J. MAnorm: a robust model for quantitative comparison of ChIP-Seq data sets. Genome Biol 13, R16, doi:10.1186/gb-2012-13-3-r16 (2012).

61. Ramirez, F. et al. deepTools2: a next generation web server for deep-sequencing data analysis. Nucleic Acids Res 44, W160–165, doi:10.1093/nar/gkw257 (2016).

62. Thorvaldsdóttir, H., Robinson, J. T. & Mesirov, J. P. Integrative Genomics Viewer (IGV): high-performance genomics data visualization and exploration. Briefings in bioinformatics 14, 178–192 (2013).

63. Heinz, S. et al. Simple combinations of lineage-determining transcription factors prime cis-regulatory elements required for macrophage and B cell identities. Molecular cell 38, 576–589 (2010).

64. McLean, C. Y. et al. GREAT improves functional interpretation of cis-regulatory regions. Nature biotechnology 28, 495 (2010).

65. Ashburner, M. et al. Gene ontology: tool for the unification of biology. Nature genetics 25, 25–29 (2000).

66. Consortium, G. O. The gene ontology resource: 20 years and still GOing strong. Nucleic acids research 47, D330–D338 (2019).

67. Bult, C. J. et al. Mouse Genome Database (MGD) 2019. Nucleic Acids Res 47, D801–D806, doi:10.1093/nar/gky1056 (2019).

68. Wolf, F. A., Angerer, P. & Theis, F. J. SCANPY: large-scale single-cell gene expression data analysis. Genome Biol 19, 15, doi:10.1186/s13059-017-1382-0 (2018).

